# Single-cell RNA-sequencing identifies a novel subset of CD9^+^ reparative macrophages modulated by CCN2 derived from Nlrx1 deficient cardiomyocytes in the infarcted heart

**DOI:** 10.1101/2023.04.06.535957

**Authors:** Jun Li, Guangxiang He, Linyi Wang, Haijuan Wang, Ruixue Ma, Wenjie Sun, Jintao Hu, Jun Liang, Renqi Li, Xingan Wu, Yanling Yang, Jun Chen, Rongrong Liu, Jianming Pei

## Abstract

**Rationale:** Nlrx1, a pattern recognition receptor belonging to the NLR family, is highly expressed in the heart and is significantly increased after myocardial infarction in vivo, however, the role of Nlrx1 in post-infarction inflammation and healing is poorly understood. Although the effects of infiltrating macrophages on survival of cardiomyocytes and healing of injured hearts have been intensely studied, whether or not and how surviving cardiomyocytes might influence the phenotype and function of macrophages in the infarcted hearts was largely unexplored.

**Objective:** To investigate the role of Nlrx1 in post-infarction inflammation and healing and to dissect the underlying mechanism.

**Methods and results:** Single cell RNA-sequencing of 23,805 cells isolated from infarcted hearts revealed 7 clusters of infarct macrophages including CCR2^+^, Isg^+^, Mki67^+^ proliferative, and LYVE1^+^ cardiac resident macrophages, and identified a novel subset of CD9^+^Il1rn^+^ macrophages, which were characterized by high expression of *Cd9, Il1rn, Gpnmb* and *Spp1,* and downregulation of proinflammatory cytokines and chemokines. CD9^+^Il1rn^+^ macrophages were largely found in infarcted hearts of Nlrx1KO mice, and were transcriptionally unique with reparative phenotype. Loss of Nlrx1 attenuated ventricular dysfunction and adverse remodeling, and decreased post-infarction mortality. Flow cytometry analysis showed markedly increased number of CD9^+^ macrophages and higher mean fluorescence intensity of CD9 in Nlrx1KO mice. RNA-seq analysis revealed that CD9^+^ macrophages of Nlrx1KO infarcted hearts exhibited a reparative and anti-inflammatory phenotype as featured by upregulation of wound healing-associated genes including *CD9, Spp1* and *Fn1*, and downregulation of proinflammatory genes. Conditioned medium from Nlrx1 deficient cardiomyocytes upon hypoxia stimulation skewed macrophage polarization towards a reparative and inflammation-suppressed phenotype. Gain-and loss-of function of Nlrx1 combined with RNA-seq analysis identified CCN2, whose expression and secretion were significantly increased in Nlrx1 deficient cardiomyocytes upon hypoxia, as a cardiomyocytes-derived mediator responsible, at least partly, for reprograming macrophage towards a reparative phenotype. In vivo, CCN2 expression was markedly elevated in Nlrx1KO infarcted hearts. Moreover, exogenous CCN2 partially mediated macrophage transition towards a reparative phenotype.

**Conclusions:** CD9^+^ IL1rn^+^ macrophages modulated by Nlrx1 deficient cardiomyocytes-derived CCN2, is a novel subset of reparative macrophages contributing to infarct healing. Inhibition of myocardial Nlrx1 and/or activation of CCN2 to boost CD9^+^ IL1rn^+^ reparative macrophages may represent new therapeutics for infarct healing following ischemic injury.

## Introduction

Myocardial infarction (MI) results in massive cell death of ischemic myocardium causing release of damage-associated molecular patterns that stimulate the innate immune response. Monocytes/macrophages infiltration and activation are the hallmark of inflammatory response following MI, and macrophages of infarct microenvironment exhibiting subsets heterogeneity and functional plasticity are essential for healing of infarcted heart.^1,2^ Here, our study with single cell RNA-sequencing (RNA-seq) revealed heterogeneity of infarct macrophages subsets including CCR2^+^ proinflammatory macrophages, Mki67^+^ proliferative macrophages, Isg^+^ interferon-inducible macrophages and LYVE1^+^ cardiac resident macrophages. Of note, we identified for the first time a novel cluster of CD9^+^ macrophages in the infarcted hearts, which were characterized by high expression of *Cd9, Il1rn, Gpnmb, Spp1* and *Atp6v0d2* and were transcriptionally unique with reparative phenotype. CD9^+^Il1rn^+^ macrophages were largely found in infarcted hearts of Nlrx1KO mice, suggesting that the presence of CD9^+^Il1rn^+^ infarct macrophages was associated with loss of Nlrx1.

Nlrx1 is a recently identified pattern recognition receptor belonging to the nucleotide-binding-domain and leucine-rich-repeat (NLR) family. Although originally suggested to be a negative regulator of innate immune responses to viruses,^3^ accumulating evidences show that Nlrx1 exerts diverse regulatory effects on innate immune responses depending on cell context and pathogenetic or experimental conditions.^4–7^ The role of Nlrx1 in tissue injury and repair is also controversial as illustrated by Kors’s studies showing that loss of Nlrx1 results in increased oxidative stress and apoptosis in epithelial cells during renal ischemia-reperfusion injury,^8^ whereas, deletion of Nlrx1 increases fatty acid metabolism and prevents diet-induced hepatic steatosis and metabolic syndrome.^9^

Nlrx1 is suggested to be highly expressed in the heart, ^3^ and our results revealed high expression of Nlrx1 in cardiomyocytes and elevated level of Nlrx1 in the infarcted hearts, however, the specific role and functional significance of Nlrx1 in post-infarction inflammation and healing is poorly understood. Here, we found that global loss of Nlrx1 attenuated ventricular dysfunction and adverse remodeling and decreased post-infarction mortality, which was associated with increased CD9^+^IL1rn^+^ macrophages in the Nlrx1KO infarcted hearts, and CD9^+^CD11b^+^ infarct macrophages exhibited a reparative and anti-inflammatory phenotype. Given the high expression of Nlrx1 in cardiomyocytes relative to that in macrophages, we hypothesized that the presence of CD9^+^IL1rn^+^ reparative macrophages may be largely attributed to deficiency of Nlrx1 in cardiomyocytes, and certain mediator(s) secreted by Nlrx1 deficient cardiomyocytes upon ischemic injury may modulate infarct macrophages towards a reparative and/or anti-inflammatory phenotype, thereby contributing to infarct healing. Conditioned medium from Nlrx1 deficient cardiomyocytes upon hypoxia stimulation skewed macrophage polarization towards a reparative and inflammation-suppressed phenotype. Gain- and loss-of function of Nlrx1 combined with RNA-seq analysis identified CCN2, which is significantly elevated in the infarcted hearts of Nlrx1KO mice in vivo and secreted by Nlrx1 deficient cardiomyocytes upon hypoxia, as a cardiomyocytes-derived mediator responsible, at least partly, for reprograming macrophage towards a reparative phenotype. Our study establishes a new paradigm for the cross-talk between cardiomyocytes-derived mediators and macrophages of infarcted hearts.

## Methods

A detailed description of the methods is provided in the Supplementary material online.

## Results

### Nlrx1 loss mitigates ventricular dysfunction, adverse remodeling and mortality after myocardial infarction

Nlrx1, a recently identified pattern recognition receptor belonging to the NLR family, is highly expressed in the hearts,^3^ however, the role of Nlrx1 in myocardial ischemic injury and healing is poorly understood. Our in vivo results showed that Nlrx1 expression was significantly increased in the infarcted hearts 3 days after MI (Figure 1A). To determine the effect of Nlrx1 loss on ventricular function and cardiac remodeling, Nlrx1KO and wild type (WT) mice were subjected to MI by ligation of left anterior descending coronary artery. Baseline echocardiograms revealed comparable chamber dimensions and ventricular function between Nlrx1KO and WT hearts (Figure 1B-G). After 7 days of MI, both WT and Nlrx1KO hearts developed decreased LV ejection fraction (LVEF) and increased LV chamber dimension, suggesting cardiac dysfunction and LV dilatation following infarction. Of note, loss of Nlrx1 ameliorated MI-induced ventricular dysfunction and LV dilatation, as evidenced by significantly increased LVEF, decreased LV end-systolic dimension (LVESD), and reduced LV end-diastolic dimension (LVEDD) in Nlrx1KO mice compared with WT mice (Figure 1B-G). Moreover, Nlrx1KO mice exhibited significantly decreased post-MI mortality in comparison to WT mice (Figure 1H).

**Figure 1.**
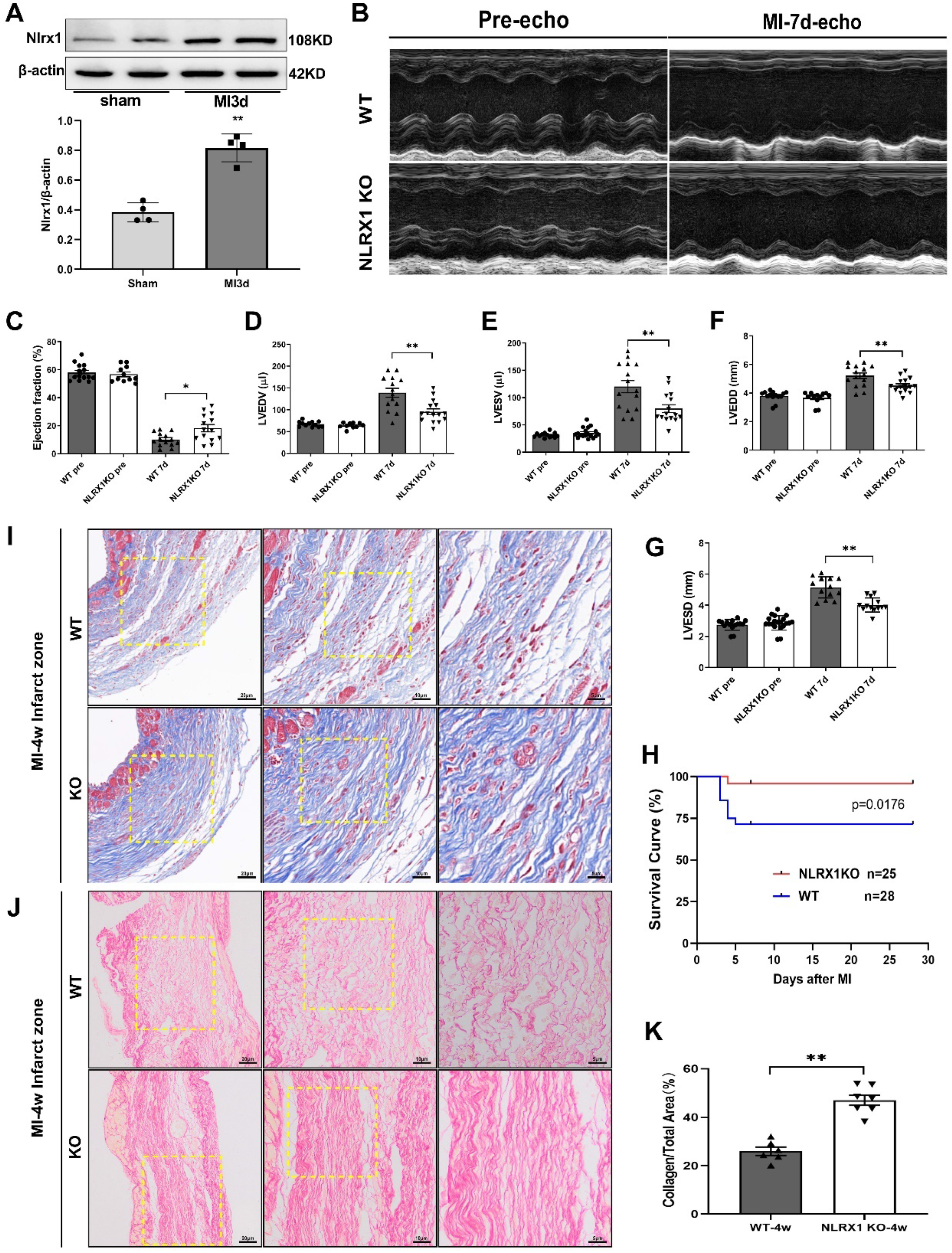
Loss of Nlrx1 mitigates ventricular dysfunction, adverse remodeling and mortality following myocardial infarction. **A**, Representative immunoblots and summary data showing increased expression of Nlrx1 in infarcted heart tissue 3 days after MI. ** *P*<0.01 versus sham, n=4 per group. **B**, Representative M-mode echocardiograms of WT and Nlrx1KO mice at baseline (pre-MI) and 7 days post-MI. **C-G**, Echocardiographic analysis of left ventricle ejection fraction (LVEF), LV end-diastolic volume (LVEDV), LV end-systolic volume (LVESV), LV end-diastolic diameter (LVEDD) and LV end-systolic diameter (LVESD) at baseline and on day 7 after MI. n=11-15 as indicated. * *P*<0.05, ** *P*<0.01. Data are expressed as mean±SEM. **H**, Kaplan-Meier analysis of survival curves between Nlrx1KO and WT mice. Post-MI mortality was significantly decreased in Nlrx1KO group. *P*=0.0176, WT, n=28; Nlrx1KO, n=25. **I-K**, Representative images of masson’s trichrome (**I**) and picrosirius red staining (**J**) showing increased collagen deposition in the infarct zone of Nlrx1KO mice 4 weeks after MI. Well-healed scar of Nlrx1KO infarct was predominated by closely assembled and well-aligned collagen fibers, in contrast to disorganized scar formation with a predominance of loosely and fragmented collagen fibers in WT mice. Scale bar as indicated. **K**, collagen density in the infarct zone after 4 weeks of MI was expressed as collagen volume fraction. n=6-7 as indicated, ** P<0.01. Data are expressed as mean±SEM.

Histological analysis revealed that there was no morphological difference of basal hearts between WT and Nlrx1KO mice. After 4 weeks of MI, Nlrx1KO mice exhibited improved infarct healing as illustrated by increased collagen deposition in the infarct region (Figure 1I-K). In addition, both masson’s trichrome and picrosirius red staining revealed well-healed scars with closely assembled and well-aligned collagen fibers in Nlrx1KO infarcts in contrast to disorganized scar formation with a predominance of loosely and fragmented collagen fibers in WT mice, suggesting reparative collagen deposition and improved infarct healing in Nlrx1KO mice (Figure 1I-J, and Supplementary Figure S1). Collectively, these results demonstrate that Nlrx1 loss ameliorates ventricular dysfunction and adverse remodeling following infarction, contributing to increased post-infarction survival of Nlrx1KO mice.

### Single-cell RNA-seq analysis reveals heterogeneity of cardiac interstitial cells following myocardial infarction

Besides cardiomyocytes, the heart contains numerous interstitial cell types including fibroblast, vascular and immune cells, which play essential roles in cardiac injury and repair. To explore the cell population heterogeneity of infarcted hearts and contribution of distinct cell types to cardiac injury and healing, we performed single-cell RNA-seq on cardiac single cells isolated from infarcted hearts (infarct zone and border zone) after 3 days of MI. Totally 23,805 single cells including 12,180 cells from 3 WT infarcted hearts and 11,625 cells from 3 Nlrx1KO infarcted hearts were sequenced, and 11 clusters with distinct lineage identities and transcriptional profiles were identified (Figure 2A-F). Interstitial cell types of infarcted hearts comprised macrophages (Cd68^+^Cd163^+^Lyz2^+^, 65.5%), fibroblasts (Col1a1^+^Col1a2^+^Dcn^+^Lum^+^; 14.3%), endothelial cells (Cdh5^+^Pecam1^+^Kdr^+^; 6.8%), monocytes (Ly6c^+^Lyz2^+^Sell^+^; 4.2%), dendritic-like cells (Cd209a^+^H2-Ab1^+^Clec10a^+^; 4.3%), neutrophils (Csf3r^+^Cxcr2^+^S100a8^+^; 2.9%), pericytes (Rgs5^+^Mcam^+^Pdgfrb^+^; 1.1%), smooth muscle cells (SMCs; Acta2^+^Tagln^+^Mylk^+^Myh11^+^; 0.4%), T-cells (Cd3d^+^Cd2^+^Trac^+^; 0.4%) and B-cells (Ms4a1^+^Cd79a^+^Cd79b^+^; 0.05%) (Figure 2A-F).

**Figure 2.**
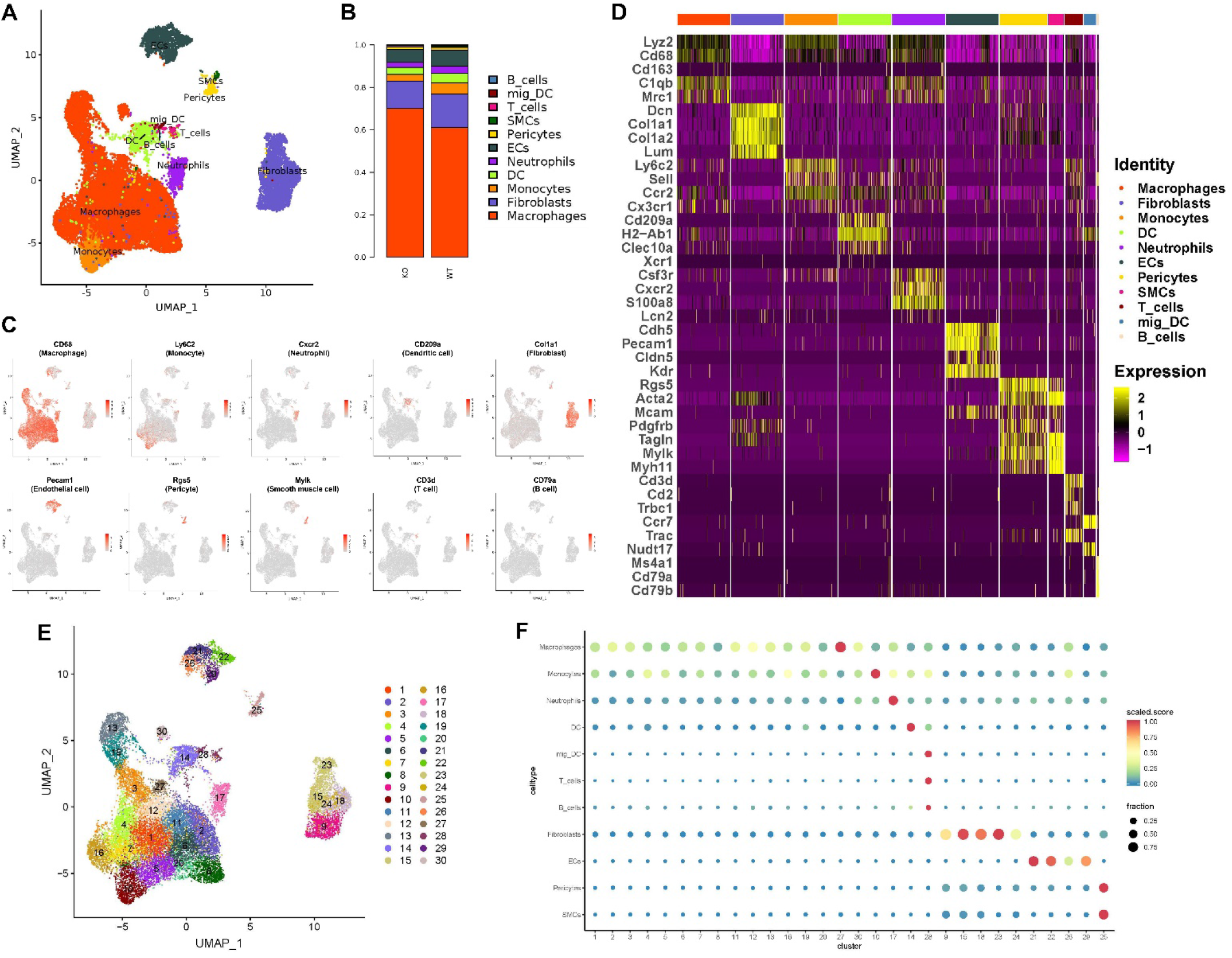
Single-cell RNA-seq analysis of cardiac interstitial cells of infarcted hearts. **A**, Single cell RNA sequencing of 23,805 cells from infarcted hearts of WT (12,180 cells from 3 pooled hearts) and Nlrx1KO (11,625 cells from 3 pooled hearts) mice clustered 11 major cell populations. Cell lineage inferred from expression of signature gene. DC, dendritic cell; EC, endothelial cell; SMC, smooth muscle cell. **B**, Cell proportion of each lineage population annotated by group. **C**, Feature plots depicting single-cell gene expression of lineage signatures. **D**, Heatmap showing top 5 differentially expressed genes in each cluster of lineages, with cluster marker genes labelled on the left (top, color-coded by cluster and lineage). **E**, Totally, 30 sub-clusters were identified in the infarcted hearts 3 days after MI. **F**, Dot plot annotating cell clusters of infarcted heart by lineage signatures. Circle size indicates cell fraction expressing signature greater than mean; color indicates mean signature expression (red, high; blue, low).

Fibroblasts accounted for 14.3% of cardiac interstitial cell sequenced (Figure 2A-B), which is consistent with previous reports that fibroblasts represent ∼15% of cardiac nonmyocytes^10^ and are proposed to have sentinel, paracrine, extracellular matrix (ECM) and mechanical functions contributing to infarct healing and remodeling.^11,12^ Within the fibroblast lineage, we identified 4 sub-clusters including fibroblasts, myofibroblasts, pro-myofibroblasts and mesothelial cells, which accounted for 48.6-54.8%, 34-36.8%, 10.8-13.9%, and 0.4-0.7% of all sequenced fibroblasts respectively. All sub-population revealed high expression of canonical fibroblast markers like Col1a1, Col1a2, Dcn and Lum, albeit at varying proportions and levels. Fibroblasts were characterized by high expression of Gsn, Pi16, Clec3b, Htra3 and Apod. Myofibroblasts were featured by higher levels of Cxcl12, Prg4, Mt2, Ptn and Isg15 than other clusters of fibroblasts. Pro-myofibroblasts showed significant expression of cell proliferation markers like Mki67, Top2a, Pclaf, Prc1 and Tpx2, which are indicative of proliferative renewal of fibroblasts in the infarcted hearts. Mesothelial cells exhibited high expression of Saa3, Clu, Dmkn, Cxadr and Ptgs1. There were no significant differences between WT and Nlrx1KO group with respects to fibroblast gene transcription and pathway analysis (Supplementary Figure S2A-D).

Endothelial cells (ECs) comprised 6.8% of cardiac single cells sequenced (Figure 2A-B), and there were three sub-populations of ECs including vascular, proliferative and lymphatic ECs, which accounted for 76.6-79.4%, 18-21.1%, and 2.4% of all sequenced ECs respectively. Vascular ECs expressed canonical endothelial markers including Cdh5, Pecam1, Cldn5 and Kdr, and were comparable in number and percentage between WT and Nlrx1KO mice. Proliferative ECs were characterized by high expression of Mki67, Top2a and Prc1, suggesting cycling of endothelial cells in the healing hearts. Lymphatic lineage ECs almost uniquely expressed Mmrn1, Lyve1 and Prox1, consistent with a lymphatic identity. There were no significant differences between WT and Nlrx1KO group regarding gene transcription and pathway analysis (Supplementary Figure S3A-D). Moreover, our EC data are broadly consistent with recently published single-cell data in the setting of infarcted hearts.^13^

Monocytes comprised 4.2% of cardiac interstitial cells (Figure 2A-B), and two sub-clusters including classical monocytes and nonclassical monocytes were identified. Classical monocytes, accounting for 96.9% of monocytes sequenced, expressed canonical monocyte markers including Ly6c, Lyz2 and Sell, and cluster specific genes like Ccl2, C3ar1, Fn1 and Ccr1. Nonclassical monocytes, accounting for 3.1% of monocytes, showed high expression of Ear2, Adgre4 and Ceacam1. Percentage of monocytes subsets, gene transcription and pathway analysis were comparable between WT and Nlrx1KO group (Supplementary Figure S4A-D). Dendritic cells (DCs) accounted for 4% of cardiac cells, and comprised two subsets of conventional DCs, namely cDC1 and cDC2, accounting for 10% and 90% of DCs respectively. cDC2 expressed Mgl2 and Ctsd, as well as typical DC marker Cd209a, and likely represents canonical DCs, whereas cDC1 highly expressed Naaa, Irf8 and Ifi205. However, there were no significant differences between WT and Nlrx1KO group regarding percentage of DC subsets, gene transcription and pathway analysis (Supplementary Figure S5A-D).

Neutrophils comprised 2.9% of cardiac single cells sequenced (Figure 2A-B), and 6 sub-clusters were identified. All sub-population revealed high expression of canonical neutrophil markers including Csf3r, Cxcr2 and S100a8. Percentage of neutrophil subsets, gene transcriptio and pathway analysis were comparable between WT and Nlrx1KO group, albeit at varying proportions and levels (Supplementary Figure S6A-D). Additionally, pericytes (Rgs5^+^Mcam^+^Pdgfrb^+^), smooth muscle cells (Acta2^+^Tagln^+^Mylk^+^Myh11^+^), T-cells (Cd3d^+^Cd2^+^Trac^+^) and B-cells (Ms4a1^+^Cd79a^+^Cd79b^+^) were also identified in the infarcted hearts, though accounting for less than 1% of all cardiac interstitial cells.

### Single cell RNA-seq analysis reveals heterogeneity and diversity of macrophages in the infarcted hearts

As the most abundant cell types in the infarcted hearts, macrophages accounted for approximately 65% of all cardiac single cells sequenced and were divided into 7 clusters exhibiting a broad range of phenotypic heterogeneity and functional diversity (Figure 3A–B). Cluster 1 of macrophages accounted for 51.3% of infarct macrophages following 3 days of MI, and expressed canonical macrophage marker genes including *Cd68, Lyz2* and *adgre1*, as well as *Arg1, Hmox1* and *pf4* (Figure 3B–E; Supplementary Figure S8A). The number and percentage of cluster 1 macrophage were comparable between WT and Nlrx1KO mice (Figure 3B–E).

**Figure 3.**
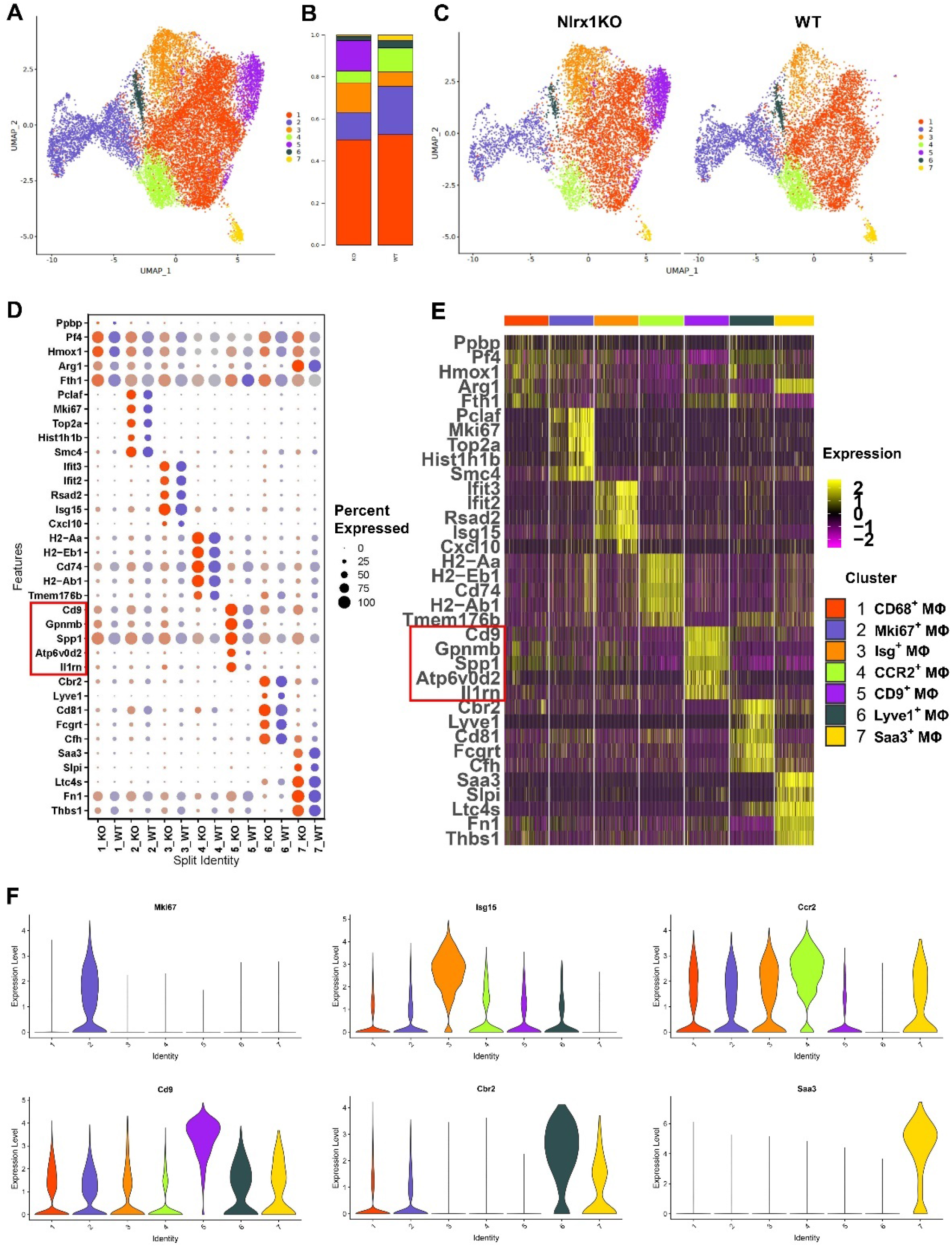
Single-cell RNA sequencing reveals heterogeneity and diversity of cardiac macrophages in the infarcted hearts. **A**, UMAP plot depicting 7 clusters of macrophages in the infarcted hearts 3 days after MI. **B**, Fraction of each cluster of macrophages in WT and Nlrx1KO group. Cluster 5 (pink) accounted for 14.6% of infarct macrophages in Nlrx1KO mice, relative to that of 0.2% in WT mice. **C**, UMAP plot showing macrophage clusters annotated by group. Cluster 5 of macrophages (pink) was mainly detected in Nlrx1KO infarcted hearts. **D**, Dot plot of top 5 upregulated genes of each macrophage cluster across conditions (red denote Nlrx1KO; blue denote WT). Circle size indicates cell fraction expressing signature gene. **E**, Heatmap showing top 5 differentially expressed genes of each macrophage cluster after 3 days of infarction (top, color-coded by cluster). **F**, Violin plots depicting differential expression of cluster-defining marker genes across subsets of infarct macrophages.

Cluster 2 of macrophages significantly expressed proliferation-indicative genes like *Pclaf, Mki67, Smc4* and *Top2a*, and was identified as proliferating macrophage (termed Mki67 cluster), accounting for 13%-23% of the infarct macrophages (Figure 3B–F; Supplementary Figure S8B). Infarcted hearts of WT mice showed more Mki67^+^ cycling macrophages relative to Nlrx1KO mice, highlighting the potential role of proliferative expansion in promoting pro-inflammatory macrophages accumulation early after infarction.

Cluster 3 of macrophages were featured by high expression of interferon-stimulated genes such as *Isg15, Ifit3, Ifit2* and *Rsad2*, and was termed as Isg cluster,^14^ accounting for around 10.6% of infarct macrophages (14% in Nlrx1KO hearts vs 6.9% in WT hearts) (Figure 3B–F; Supplementary Figure S7C, S8C).

Cluster 4 of macrophages exhibited high expression of major histocompatibility complex (MHC)-II associated genes (such as *H2-Eb1, H2-Aa1, H2-Ab1*) and typical proinflammatory genes like *Ccr2*, *Il1b*, *Plbd1* and *Tmem176b*, and was identified as MHC-II^+^CCR2^+^ macrophages (Figure 3C–D; Supplementary Figure S7B, S8D). Notably, the number and percentage of MHC-II^+^CCR2^+^ macrophages were significantly reduced in Nlrx1KO infarcts compared with WT (5.5% in Nlrx1KO hearts vs 11.2% in WT hearts) (Figure 3B–F), suggesting suppression of proinflammatory CCR2^+^ macrophages in Nlrx1KO infarcted hearts.

Cluster 5 of macrophages were characterized by high expression of *Cd9, Il1rn, Gpnmb, Spp1* and *Atp6v0d2*, and was termed as CD9 cluster, which was not reported to date in the infarcted hearts. Intriguingly, CD9^+^Il1rn^+^ macrophages accounted for 14.6% of infarct macrophages in Nlrx1 KO mice, relative to that of 0.2% in WT mice, suggesting that the presence of CD9^+^Il1rn^+^ macrophages in the infarcted hearts was associated with loss of Nlrx1 (Figure 3B–F; Supplementary Figure S7A).

Cluster 6 of macrophages were enriched in *Cbr2, Lyve1, Fcgrt* and *Timd4*, and was termed as LYVE1^+^ cardiac resident macrophages accounting for 2.7% of infarct macrophages (Figure 3B–F; Supplementary Figure S7D, S8E), which is consistent with previous finding that LYVE1^+^Timd4^+^MHC-II^low^CCR2^-^ macrophages are cardiac tissue-resident population contributing to homeostasis via phagocytosis of dead cardiomyocytes.^15^

Cluster 7 of macrophages showed high expression of *Saa3, Slpi, Ltc4s* and *Thbs1* and comprised the minority (1.7%) population of the infarct macrophages (Figure 3B–F; Supplementary Figure S8F).

### CD9^+^IL1rn^+^ macrophages exhibit unique transcriptional profile and functional characteristics

As a novel cluster of macrophages identified in this study, CD9^+^Il1rn^+^ macrophage was not reported to date in the infarcted hearts by other studies using single-cell RNA-seq or other approaches. In order to delineate the transcriptional profile of CD9^+^Il1rn^+^ macrophages, differentially expressed genes and classical marker genes associated with macrophage phenotype and activation were analyzed. Wound healing and inflammation-resolution associated genes including *Cd9, Gpnmb, Spp1, Atp6v0d2* and *Il1rn* were upregulated, whereas proinflammatory cytokines (*Il1b*) and chemokines (*Ccr2, Ccl2, Ccl4, Ccl7, Ccl12*),^16^ and *Plbd1* (an independent marker of cardiac dysfunction post-MI)^17^ were significantly downregulated in CD9^+^Il1rn^+^ macrophages (Figure 4A; Supplementary Figure S9-S10).

**Figure 4.**
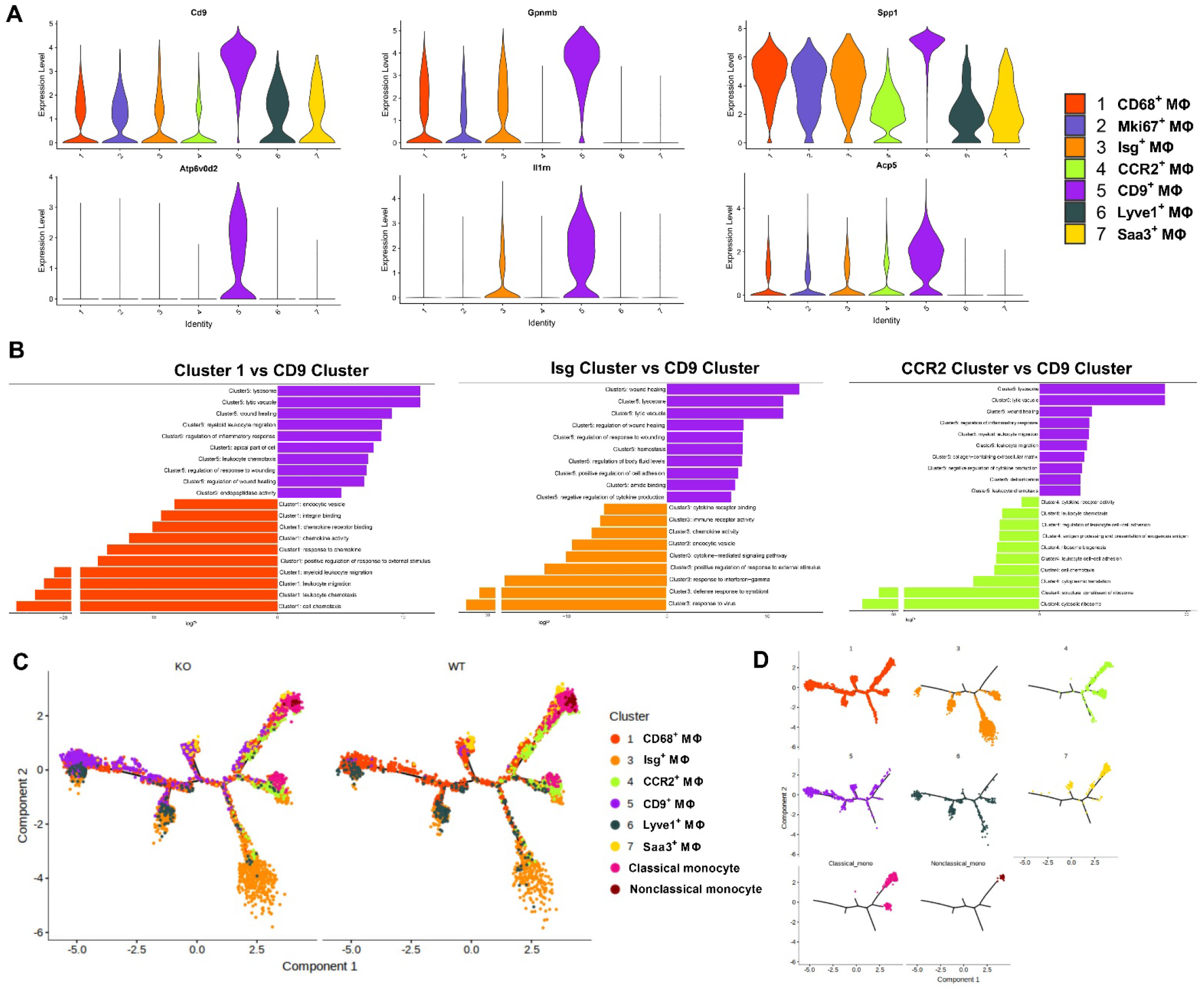
Transcriptional profile and functional characteristics of CD9^+^IL1rn^+^ macrophages. **A**, Violin plots depicting differential expression of CD9 cluster-defining genes across 7 subsets of infarct macrophages. CD9 cluster exhibited high expression of wound healing and inflammation resolution associated genes including *Cd9, Spp1, Il1rn, Gpnmb* and *Atp6v0d2* relative to other clusters. **B**, Pathway enrichment analysis of differentially expressed genes in CD9 cluster versus other clusters of macrophages. Pathway enrichment is expressed as the –log[P] adjusted for multiple comparison. **C**, Monocle trajectory analysis annotated by group. Different clusters of macrophages occupy distinct trajectory branches with somewhat overlapping, indicative of distinct states of differentiation and activation. **D**, Monocle trajectory analysis shows distinct ontogeny of CD9 cluster, which is different from that of CCR2 cluster and ISG cluster with less overlapping.

Then, we performed pathway analysis by individually comparing CD9 cluster to other clusters. Upregulated pathways of CD9 cluster were enriched in homeostatic and reparative functions including endocytosis, lysosome function, wound healing, regulation of wound healing, negative regulation of cytokine production and inflammatory response, and detoxification. Whereas classic inflammatory pathways, including leukocytes chemotaxis, leukocyte cell-cell adhesion, cytokine receptor activity, antigen processing and presentation, and translational-ribosomal pathways (cytosolic ribosome, and cytoplasmic translation) were remarkably enriched in CCR2 cluster relative to CD9 cluster (Figure 4B; Supplementary Figure S11-S12). Pathways relating defense response to virus and external stimulus, and interferon-gamma responsive pathways were enriched in Isg cluster. Pathways associated with leukocytes chemotaxis, leukocytes migration, response to chemokine, and chemokine activity were upregulated in Arg1 & Hmox1 cluster, whereas proliferation-related pathways including chromosome segregation, nuclear division, organelle fission, were upregulated in Mki67 cluster (Figure 4B; Supplementary Figure S11-S12). These data reveal the unique and cluster-specific functions of macrophages in the infarcted hearts.

To determine the ontogeny of CD9^+^ macrophages and developmental relationships with other monocytes/macrophage clusters, pseudotime trajectory analysis by the Monocle algorithm revealed that distinct clusters of macrophages occupied different trajectory branches with more or less overlapping (Figure 4C-D). Cluster 5 of CD9^+^ macrophages shared part of trajectory branches occupied by cluster 1, which straddled almost all trajectory branches. In contrast to CD9 cluster, CCR2 cluster occupied distinct trajectory branches and largely overlapped with monocytes clusters, suggesting the ontogeny of CCR2^+^ macrophages were closely associated with monocytes, which is consistent with previous finding that proinflammatory CCR2^+^ macrophages derived from circulating monocytes infiltrating the infarcted heart.^18,19^ Isg cluster occupied different trajectory branches relative to other clusters (Figure 4C-D). Of note, the trajectory branches occupied by CD9 cluster was different from that of CCR2 cluster and ISG clusters with less overlapping, suggesting distinct ontogeny of CD9 cluster, which may account partly for the unique transcriptional characteristics of CD9 cluster relative to other macrophage subsets.

Overall, these data suggest that cardiac CD9^+^ macrophages are distinct from other clusters and represent a unique subset of macrophages with anti-inflammatory and reparative function within the infarct healing niche. In contrast to other clusters including CCR2^+^ and Isg^+^ macrophages, which have been reported in previous studies,^14,18,19^ CD9^+^ macrophage was identified, for the first time to our knowledge, in the setting of infarcted hearts.

### CD9^+^ macrophages of Nlrx1KO infarcted hearts exhibit a reparative and anti-inflammatory phenotype

To further confirm the findings of single cell RNA-seq study, we analyzed cardiac macrophages isolated from infarcted hearts by flow cytometry. The number and percentage of cardiac CD11b^+^Ly6G^-^ cells were comparable between WT and Nlrx1KO mice, however, the number of CD9^+^CD11b^+^Ly6G^-^ macrophages were markedly increased in Nlrx1KO infarcted hearts (72.2% in Nlrx1KO vs 40.9% in WT) (Figure 5A-B), which was consistent with our single cell RNA-seq finding that CD9^+^ macrophages were largely present in Nlrx1KO hearts (Fig3B-C). Moreover, the mean fluorescence intensity of CD9 of Nlrx1KO infarct macrophages was significantly higher than that in WT group (227 in Nlrx1KO vs 85.3 in WT) (Figure 5C).

**Figure 5.**
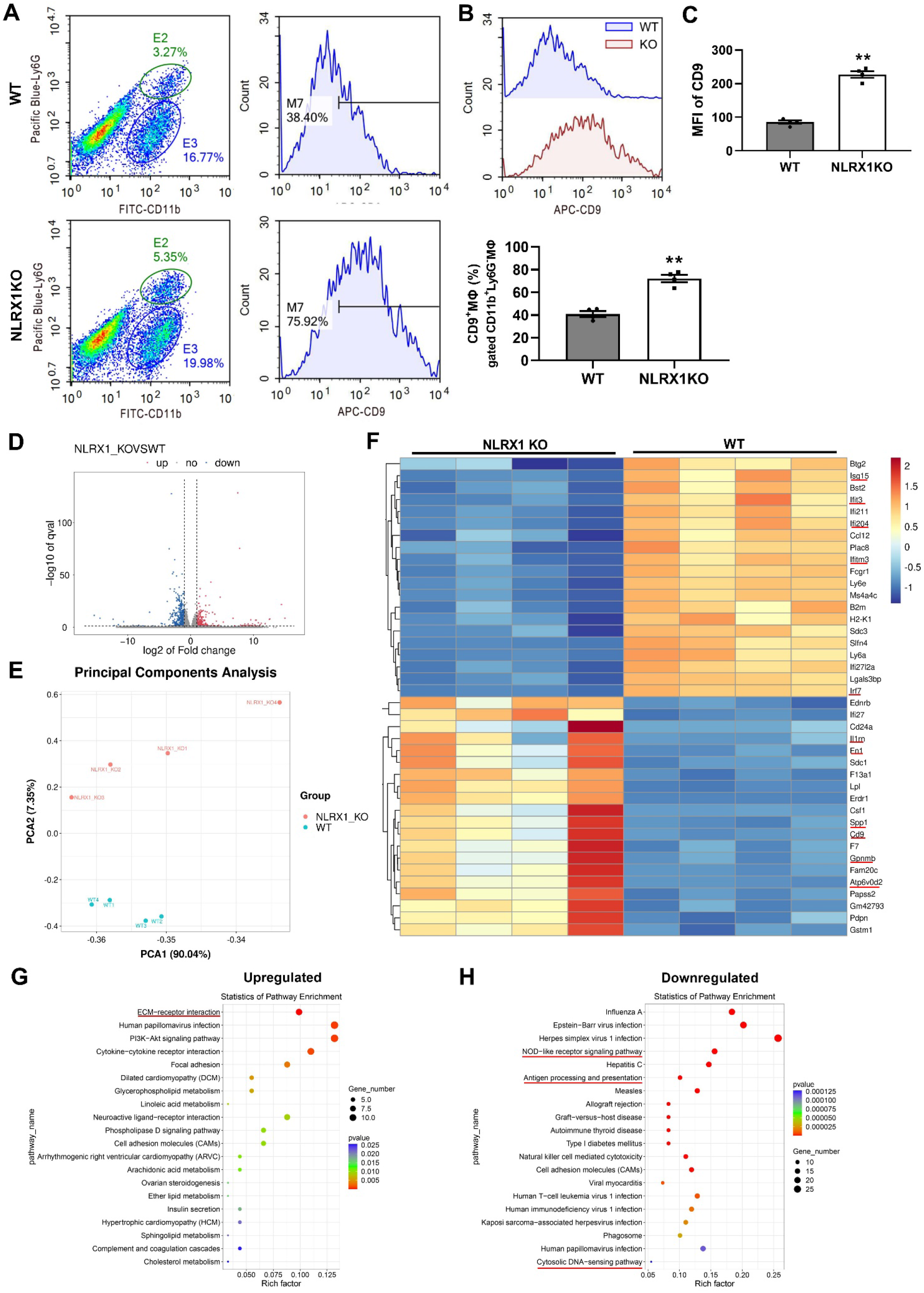
CD9^+^CD11b^+^ macrophages of infarcted hearts exhibit a reparative and anti-inflammatory phenotype. **A-B**, Flow cytometry analysis showing a marked increase of CD9^+^CD11b^+^Ly6G^-^ macrophages in Nlrx1KO infarcted hearts, though comparable number and percentage of CD11b^+^Ly6G^-^ cells between Nlrx1KO and WT mice. **C**, Mean fluorescence intensity of CD9 of Nlrx1KO infarct macrophages was significantly higher than that of WT group. n=4, ** *P*<0.01 vs WT. **D**, Volcano plot illustrating differentially expressed genes (DEGs) in CD9^+^CD11b^+^ macrophages between Nlrx1KO and WT mice, based on >2-fold and with adjusted P value (padj) <0.05. **E**, Principal components analysis of CD9^+^CD11b^+^ infarct macrophages. The distance of the dots indicates the similarity of transcriptome: the greater difference between the transcriptomes of macrophages, the greater distance between the dots. **F**, Heatmap showing top 20 DEGs with FPKM (fragments per kilobase of transcript sequence per millions base pairs sequenced) value greater than 10 between Nlrx1KO and WT CD9^+^CD11b^+^ macrophages. n=4 for each condition. Inflammatory genes like Isg15, Ifit3, Ifitm3, and Irf7 were suppressed, while inflammation-resolution and wound healing associated genes including CD9, Spp1, Gpnmb, Fn1 and Il1rn were upregulated in Nlrx1KO macrophages. **G-H**, Top 20 of upregulated (G) and downregulated (H) KEGG pathway enrichment of the identified DEGs between NLRX1KO and WT CD9^+^CD11b^+^ macrophages. DEGs indicates differentially expressed genes.

In order to determine the effects of Nlrx1 loss on transcriptional profile of infarct macrophages, CD11b^+^Ly6G^-^CD9^+^ macrophages were isolated from WT and Nlrx1KO infarcted hearts on day 3 post-MI by FACS sorting, then subjected to bulk RNA-seq for genome-wide transcriptomic analysis. In CD11b^+^CD9^+^ macrophages of Nlrx1KO infarcts, 232 genes were upregulated, whereas 373 genes were downregulated in comparison to WT macrophages (>2-fold, padj<0.05) (Figure 5D-F, Supplementary Figure S13). With differentially expressed genes, we performed Gene Ontology (GO) enrichment and Kyoto Encyclopedia of Genes and Genomes (KEGG) pathway functional enrichment analysis. Among the top 20 of enriched pathways, ECM-receptor interaction, cytokine-receptor interaction and focal adhesion were upregulated, whereas, NOD-like receptor signaling pathway, cytosolic DNA-sensing pathway, immune response to virus, antigen processing and presentation, cellular response to interferon-beta, and interferon responsive pathway were significantly downregulated in Nlrx1KO infarct macrophages (Figure 5G-H; Supplementary Figure S14-S15). Moreover, most of the downregulated genes (such as *Irf7, stat1, gbp2, gbp3, Oas3, Oas2, ccl12, Nod1, stat2, Il18, Oas1a, Oas1g, Ifi204, Ifi207, Irf7, zbp1, Cxcl10, Il18, Ddx58, Trex1*) were overlapped in NOD-like receptor signaling pathway, interferon responsive pathway, and cytosolic DNA-sensing pathway, indicating suppression of these pathways in Nlrx1KO infarct macrophages (Figure 5F-H; Supplementary Figure S13-S15).

Gene-specific analysis revealed that CD11b^+^Ly6G^-^CD9^+^ macrophages of Nlrx1KO infarcts exhibited a less inflammatory transcriptional signature, as evidenced by remarkable downregulation of IFN-responsive genes including *Ifit3, Ifit2, Isg15, Rsad2*, and other proinflammatory genes like *Cxcl10, Ccl12, Ccl8, Cxcl16* and *Il18*. Moreover, NOD-like receptor signaling pathway associated genes including *Irf7, Stat1, Stat2, gbp2, gbp3, Oas3, Oas2, Oas1a, Ifi204 and Ifi207*, and inflammasome activation associated transcriptional factors like *Zbp1* were also suppressed in CD11b^+^CD9^+^ macrophages of Nlrx1KO infarcts (Figure 5F; Supplementary Figure S13B). Meanwhile, CD11b^+^CD9^+^ macrophages of Nlrx1KO infarcts exhibited high expression of anti-inflammatory genes including *Il1rn, Chil3 (YM1), Ccl22, Ccl24, Cxcl14, Cebpb, Bhlhe40* and *Arg1*, and wound healing associated genes including *CD9, Spp1, Gpnmb, Sdc1, Erdr1, CD24a* and *Fn1* (Figure 5F; Supplementary Figure S13B). However, mRNA levels of *TLR2, TLR4,* and *IL-10* were comparable between 2 groups. Collectively, consistent with our single cell RNA-seq data, these data suggest that CD11b^+^CD9^+^ macrophages of Nlrx1KO infarcted hearts exhibit an anti-inflammatory and reparative phenotype, which contribute to inflammation resolution and infarct healing of Nlrx1KO hearts following MI.

### Conditioned medium from Nlrx1 deficient cardiomyocytes upon hypoxia stimulation skews macrophage transition towards a reparative and inflammation-suppressed phenotype

We further explored the underlying mechanism responsible for the presence of CD9^+^CD11b^+^ macrophages in the Nlrx1KO infarcted hearts. Mining of The Human Protein Atlas (www.proteinatlas.org) revealed high expression of Nlrx1 in the hearts and cardiomyocytes (Supplementary Figure S16A-C), which is consistent with previous reports that Nlrx1 is highly expressed in heart tissues.^3^ To confirm this, we determined the protein expression of Nlrx1 in cardiomyocytes and macrophages by western blotting, and indeed, Nlrx1 expression was remarkably high in primary cardiomyocytes and H9C2 myocyte cell line, followed by RAW264.7 macrophages cell line, and lowest in bone marrow derived macrophages (Supplementary Figure S16D). These data suggest that the presence of CD9^+^IL1rn^+^ macrophage in the Nlrx1KO infarcted hearts may be largely attributed to deficiency of Nlrx1 in cardiomyocytes rather than macrophages, although the potential effects of Nlrx1 loss in other cell types can’t be excluded. Based on these data, we hypothesized that certain mediator(s) secreted by Nlrx1 deficient cardiomyocytes upon ischemic injury, may modulate and reprogram infarct macrophages towards a reparative and/or anti-inflammatory phenotype, thereby contributing to infarct healing, preserved cardiac function and increased survival after MI in Nlrx1KO mice.

To test this hypothesis, conditioned medium from neonatal rat ventricular myocytes (NRVMs) upon hypoxia stimulation and glucose deprivation, were employed to stimulate bone marrow derived macrophages (BMDMs), and gene expression profile was determined. As expected, mRNA expression of *CD9, Il1rn, Fn1* and *Spp1* were significantly upregulated in BMDMs stimulated by supernatants from siRNA–mediated Nlrx1 knockdown cardiomyocytes, when compared to supernatants from scrRNA transfected cardiomyocytes (Figure 6A). On the other hand, mRNA expression of inflammatory genes including *Isg15, Ifit3, Rsad2, Ifitm3, Ifi209, Cxcl10, Irf3, Irf7, Zbp1, Oas1a*, *Oasl2* and *Ms4a4c* were remarkably suppressed in BMDMs stimulated by supernatants from Nlrx1-siRNA cardiomyocytes (Figure 6B). Of note, these results are highly consistent with our in vivo findings that infarct macrophages of Nlrx1KO mice exhibited remarkable downregulation of inflammatory signature genes, but upregulation of anti-inflammatory or reparative genes (Figure 5F).

**Figure 6.**
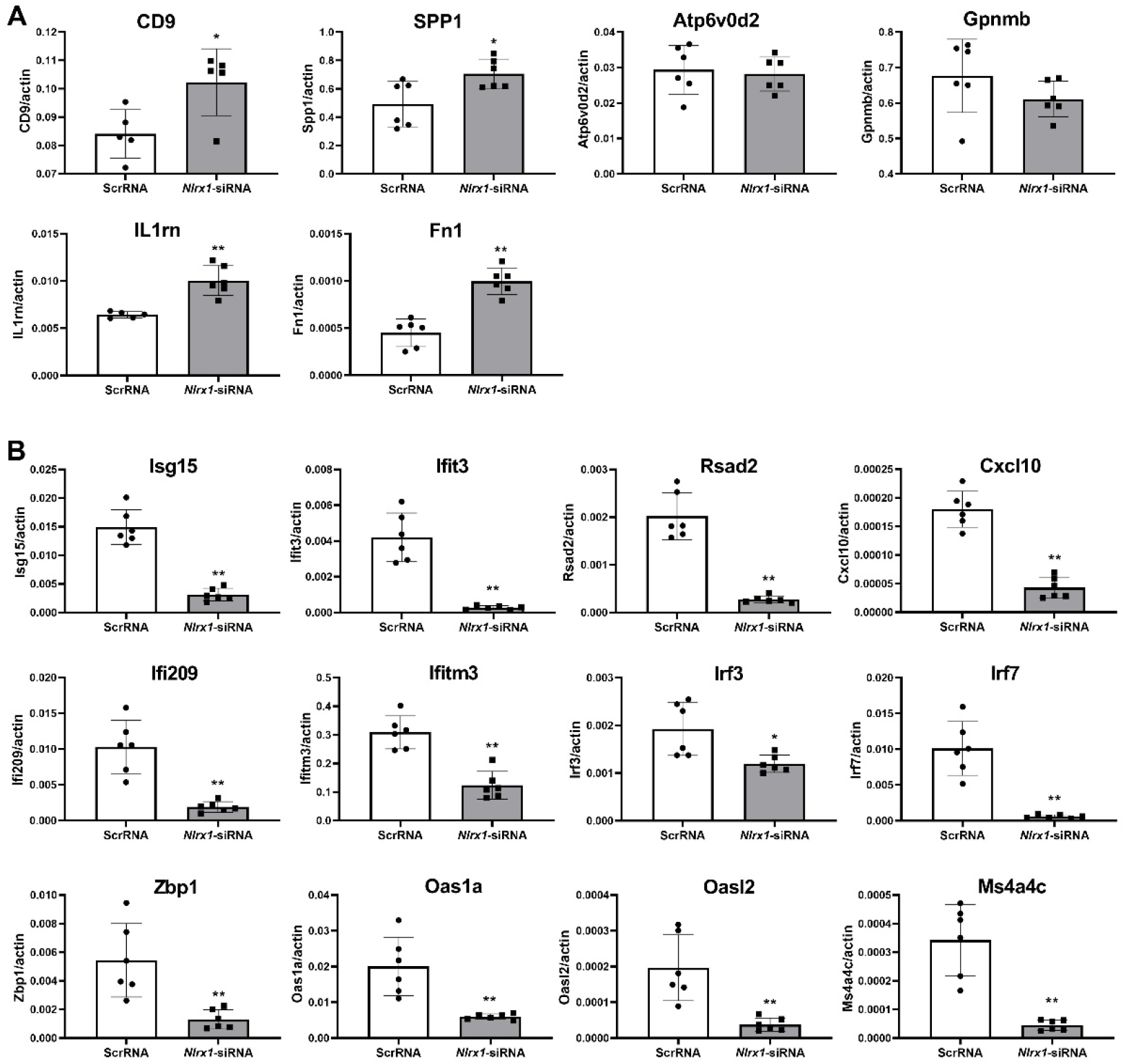
Conditioned medium of Nlrx1 deficient cardiomyocytes skews macrophage transition toward a reparative and less inflammatory phenotype. Conditioned medium from neonatal rat ventricular myocytes (NRVMs) upon hypoxia stimulation and glucose deprivation for 24 hours, were employed to stimulate bone marrow derived macrophages (BMDMs) for 24 hours. **A**, mRNA level of *CD9, Il1rn, Fn1, Spp1* were significantly upregulated in BMDMs stimulated by supernatants of Nlrx1-knockdown cardiomyocytes transfected with Nlrx1-siRNA. n=6, \**P*<0.05, \*\**P*<0.01 vs scrRNA. **B**, mRNA expression of inflammatory genes including *Isg15, Ifit3, Rsad2, Ifitm3, Ifi209, Cxcl10, Irf3, Irf7* and *Oas1a*, were remarkably suppressed in BMDMs stimulated by supernatants of Nlrx1-knockdown cardiomyocytes, compared with supernatants from scramble-siRNA transfected cardiomyocytes. n=6, \**P*<0.05, \*\**P*<0.01 vs. scrRNA.

### CCN2 is identified to be the mediator secreted by Nlrx1 deficient cardiomyocytes upon hypoxia in vitro, and increased in infarcted hearts of Nlrx1KO mice in vivo

To further identify the specific secreted mediator in the supernatants of Nlrx1 deficiency cardiomyocytes, which are responsible for modulating macrophages transition to reparative phenotype, gain-and loss-of function of Nlrx1 was carried out in NRVMs. We observed a significant knockdown of Nlrx1 by delivery of Nlrx1-targeting small interfering RNA (Supplementary Figure S17A), Then NRVMs were challenged by hypoxia stimulation and glucose deprivation for 8 hours, followed by RNA-seq analysis. Regardless of Nlrx1-siRNA or scrRNA delivery, around 2300 genes were differentially expressed in NRVMs upon hypoxia stimulation and glucose deprivation relative to normoxia control, suggesting a profound effect of hypoxia on transcriptomics of cardiomyocytes. Of note, 65 genes were upregulated and 69 genes were downregulated in NRVMs transfected with Nlrx1-siRNA in comparison to scrRNA control (>2-fold, padj<0.05) (Figure 7A-B; Supplementary Figure S17B, S18A and D). GSEA enrichment and KEGG pathway analysis of differentially expressed genes revealed that extracellular matrix (ECM) and ECM-receptor interaction ranked top of the enriched pathways in Nlrx1 knockdown NRVMs upon hypoxia versus scrRNA control. Furthermore, CCN2, as the core enriched gene of ECM set, was significantly upregulated in Nlrx1-siRNA cardiomyocytes upon hypoxia (Figure 7A and E; Supplementary Figure S17C-D, S18).

**Figure 7.**
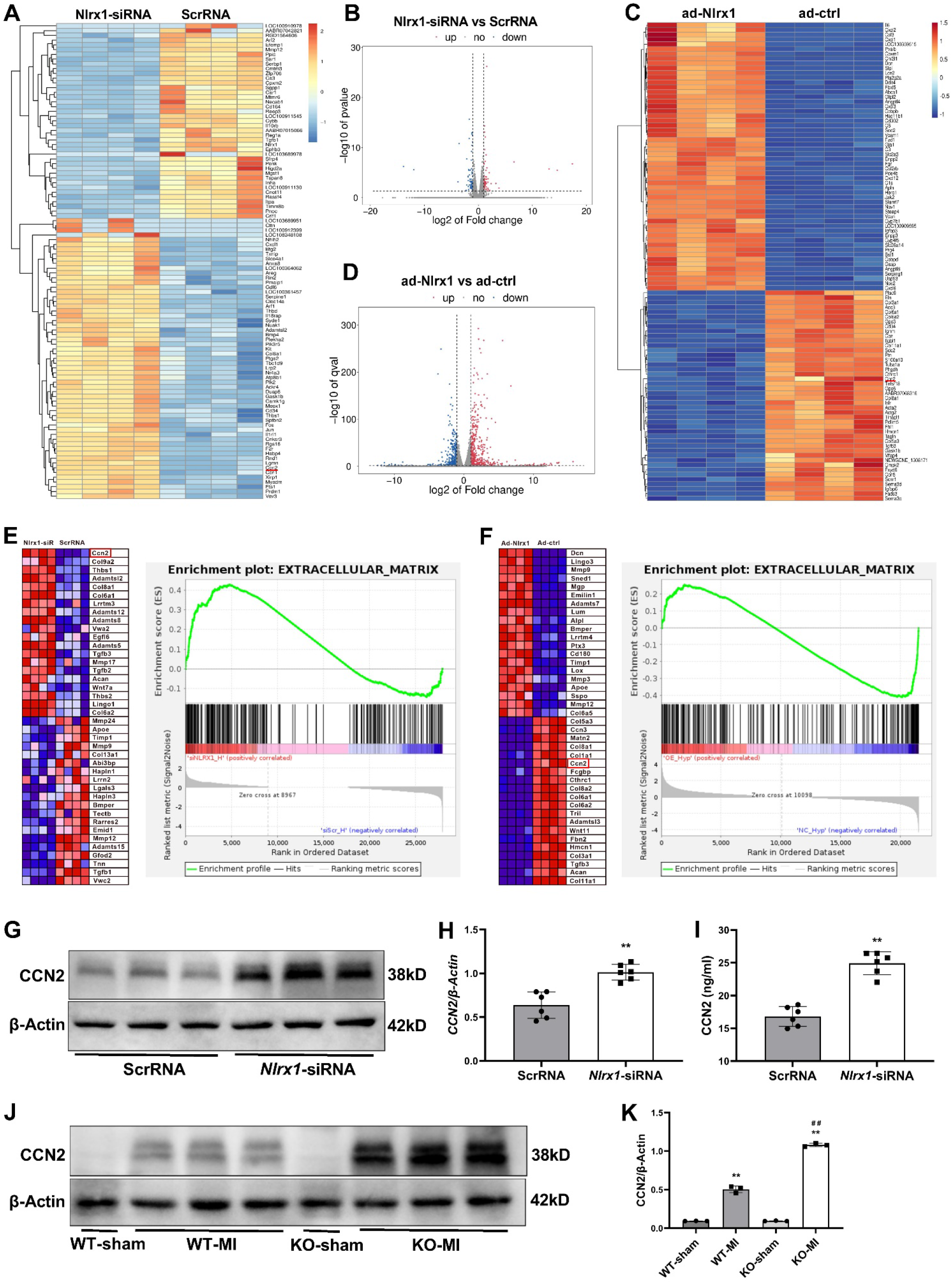
CCN2 was upregulated in NLRX1 deficient cardiomyocytes upon hypoxia stimulation in vitro, and increased in infarcted hearts of NLRX1KO mice in vivo. **A-B**, NRVMs were transfected with either Nlrx1-siRNA or scrRNA for 48 hours, then stimulated by hypoxia and glucose deprivation for 8 hours. Heat map and volcano plot illustrating DEGs of transcriptomic comparisons between Nlrx1-siRNA and scrRNA transfected NRVMs. DEGs indicates differentially expressed genes; NRVMs, neonatal rat ventricular myocytes. scrRNA, scramble-siRNA. **C-D**, NRVMs were transfected with either ad-Nlrx1 or ad-ctrl for 48 hours, then stimulated by hypoxia and glucose deprivation for 8 hours. Heat map and volcano plot illustrating DEGs between ad-Nlrx1 and ad-ctrl transfected NRVMs. ad-Nlrx1 indicates adenovirus-Nlrx1; ad-ctrl, adenovirus-vector. **E**, Top 20 core enriched genes of the extracellular matrix pathway between Nlrx1-siRNA and scrRNA transfected NRVMs upon hypoxia stimulation and glucose deprivation (left panel). GSEA analysis of extracellular matrix gene set enrichment (right panel). **F**, Top 20 core enriched genes of the extracellular matrix pathway between ad-Nlrx1 and ad-ctrl transfected NRVMs upon hypoxia stimulation and glucose deprivation. GSEA analysis of extracellular matrix gene set, NES = 2.02, *P* < 0.0001. **G**-**H**, CCN2 protein expression was remarkably increased in Nlrx1-knockdown NRVMs upon hypoxia stimulation and glucose deprivation for 24 hours. n=6, \*\**P*<0.01 vs ScrRNA. **I**, Secreted CCN2 determined by ELISA, was significantly elevated in supernatants of Nlrx1-knockdown NRVMs following hypoxia stimulation, relative to scrRNA group. n=6, \*\**P*<0.01 vs ScrRNA. **J**, Representative immunoblot showing CCN2 expression in hearts with or without myocardial infarction. **K**, CCN2 protein expression was significantly increased in the infarcted hearts of Nlrx1KO mice relative to WT control mice after 3 days of infarction. n=6, \*\**P*<0.01 vs sham, ##*P*<0.01 vs WT-MI.

NRVMs were transfected with Nlrx1-targeting adeno virus of different MOI for 48 hours, and the efficacy of Nlrx1 overexpression was confirmed by Western blotting (Supplementary Figure S19A). Then NRVMs were treated by hypoxia and glucose deprivation for 8 hours, followed by bulk RNA-seq analysis. Regardless of adeno virus-Nlrx1 (ad-Nlrx1) transfection or not, around 2555-2887 genes were upregulated and 2345-2616 genes were downregulated in NRVMs upon hypoxia and glucose deprivation relative to normoxia control, suggesting a profound change of cardiomyocyte transcriptomics following stress of hypoxia. Of note, 666 genes were upregulated and 445 genes were downregulated in NRVMs transfected with ad-Nlrx1 in comparison to ad-control (>2-fold, padj<0.05) (Figure 7C-D; Supplementary Figure S19B, S20A and D). GO and GSEA enrichment analysis of differentially expressed genes revealed that extracellular region and ECM ranked top of the enriched pathways in Nlrx1-overexpressing NRVMs upon hypoxia versus ad-control. In contrast to Nlrx1 knockdown NRVMs, Nlrx1-overexpressing NRVMs exhibited significant downregulation of CCN2 in response to hypoxia (Figure 7C and F; Supplementary Figure S19C-D, S20). Collectively, gain-and loss-of Nlrx1 study combined with unbiased RNA-seq analysis revealed that CCN2 expression was negatively correlated with the level of Nlrx1 in cardiomyocytes upon hypoxia. CCN2 was significantly upregulated in NLRX1 knockdown NRVMs, whereas, CCN2 was remarkably downregulated in NLRX1-overexpressing NRVMs in response to hypoxia stimulation and glucose deprivation.

Validation experiments revealed a significant increase of CCN2 protein expression in Nlrx1 knockdown cardiomyocytes upon hypoxia stimulation and glucose deprivation (Figure 7G-H). In addition, the secreted CCN2 in the supernatant, as determined by ELISA, was also significantly elevated in Nlrx1 knockdown cardiomyocytes relative to scramble-siRNA transfected NRVMs (Figure 7I). Collectively, both protein expression of CCN2 and secreted CCN2 in the supernatant were increased in Nlrx1 knockdown cardiomyocytes upon hypoxia stimulation. Moreover, in vivo study also demonstrated that protein expression of CCN2 was significantly elevated in the infarcted hearts of Nlrx1KO mice relative to WT mice on day 3 after infarction (Figure 7J-K). Taken together, CCN2 is significantly elevated in Nlrx1KO infarcted hearts, and CCN2 is identified to be a Nlrx1 deficient cardiomyocytes-derived mediator, responsible for reprograming macrophages towards a reparative and anti-inflammatory phenotype.

### Exogenous CCN2 mediates macrophage activation towards a reparative phenotype

We therefore focused on the effects of CCN2 on macrophage activation, and further explored whether or not exogenous CCN2 directly mediates macrophage transition to reparative phenotype. BMDMs were stimulated with supernatants from hypoxia stressed cardiomyocytes with or without recombinant CCN2 for 24 hours. Exogenous CCN2 treatment alone upregulated mRNA level of *CD9* and *Spp1* in BMDMs compared to conditioned medium of scramble-siRNA transfected cardiomyocytes (Figure 8A-B). Although *Il1rn* expression in BMDMs was not changed by exogenous CCN2 treatment alone, CCN2 addition into the conditioned medium of scramble-siRNA transfected cardiomyocytes significantly increased *Il1rn* expression (Figure 8C). These data suggest that CCN2 is responsible, at least partially, for the transition of infarct macrophages to the reparative phenotype, although the effects of other mediators derived from ischemic cardiomyocytes cannot be ruled out.

**Figure 8.**
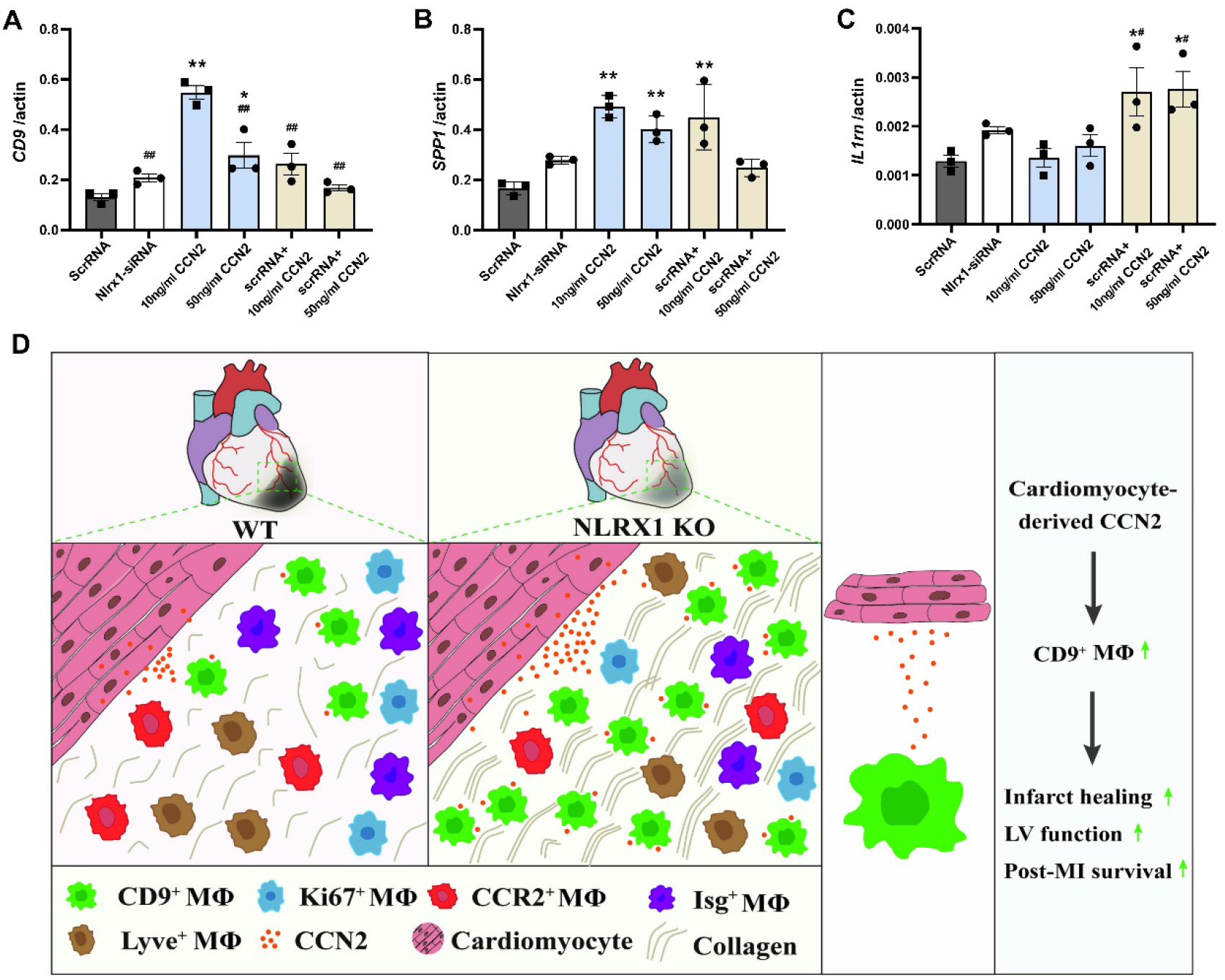
Exogenous CCN2 mediates macrophage transition to a reparative phenotype. BMDMs were stimulated with exogenous CCN2 with or without conditioned medium from neonatal rat ventricular myocytes (NRVMs) upon hypoxia stimulation and glucose deprivation for 24 hours. **A-B**, exogenous CCN2 treatment alone upregulated mRNA level of *CD9* and *Spp1* in BMDMs compared to conditioned medium of scramble-siRNA transfected cardiomyocytes. **C**, *Il1rn* expression in BMDMs was not changed by exogenous CCN2 treatment alone, however, CCN2 addition into the conditioned medium of scramble-siRNA transfected cardiomyocytes significantly increased *Il1rn* expression, suggesting CCN2 is the key mediator, in the supernatants of NLRX1 deficient cardiomyocytes, responsible for its effect on macrophage activation. n=3, \**P*<0.05, \*\**P*<0.01 vs. ScrRNA; **#***P*<0.05, **##***P*<0.01 vs. 10mg/ml CCN2. **D,** A proposed model showing the heterogeneity of macrophages in the infarcted heart, and a novel subset of reparative CD9^+^ macrophages modulated by CCN2 derived from Nlrx1 deficient cardiomyocytes, highlighting the cross-talk between cardiomyocytes-derived mediator(s) and phenotypic plasticity of cardiac macrophages. Nlrx1KO hearts exhibite increased CCN2 and more CD9^+^ reparative macrophages contributing to improved infarct healing, preserved cardiac function and decreased post-infarction mortality.

## Discussion

Our study with single cell RNA-seq analysis identified, for the first time to our knowledge, a novel cluster of CD9^+^Il1rn^+^ macrophages characterized by high expression of *Cd9, Il1rn, Gpnmb, Spp1* and *Atp6v0d2* in the infarcted hearts. CD9^+^Il1rn^+^ macrophages are largely found in Nlrx1KO hearts and are associated with improved infarct healing, preserved cardiac function and decreased post-infarction mortality in Nlrx1KO mice. RNA-seq analysis of isolated infarct macrophages revealed that CD9^+^ macrophages exhibit a reparative and anti-inflammatory phenotype with upregulation of wound healing-associated genes and downregulation of proinflammatory genes. Our in vitro and in vivo study identified CCN2, which is significantly elevated in the Nlrx1KO infarcted hearts and secreted by Nlrx1 deficient cardiomyocytes upon hypoxia, as a cardiomyocytes-derived mediator responsible, at least partly, for reprograming macrophage towards a reparative and anti-inflammatory phenotype contributing to infarct healing.

### 1. Healing role of a novel subset of CD9^+^ macrophages in the infarcted heart

A growing number of evidences demonstrate that phenotypic heterogeneity and functional diversity of infarct macrophages are hallmark of cardiac inflammation following infarction, which has been confirmed by single cell RNA-seq analysis in our study. Here, infarct macrophages are clustered into 7 subsets including Isg^+^ macrophages,^14^ CCR2^+^ macrophages,^18,19^ Mki67^+^ proliferative macrophages,^20^ LYVE1^+^ cardiac resident macrophages^15^ and CD9^+^ macrophages. Of note, infarct macrophage clusters identified by different studies using single cell RNA-seq overlapped in major clusters, with variations in percentage and amounts of clusters depending on the infarct context and time examined. Consistent with previous findings, our results revealed that Isg cluster are cardiac interferon-inducible macrophages,^14^ CCR2 cluster exhibiting high expression of major histocompatibility complex (MHC)-II associated genes are proinflammatory macrophages leading to sustained inflammation and adverse remodeling of infarcted hearts,^18,19^ and LYVE1^+^ cluster are recently identified cardiac resident macrophages.^15^ Intriguingly, our single cell RNA-seq analysis identified a novel cluster of infarct macrophages characterized by high expression of *Cd9, Il1rn, Gpnmb, Spp1* and *Atp6v0d2*, whereas Irf7, Irf3 and interferon-stimulated genes including *Isg15, Ifit3, Ifit2* and *Rsad2* were significantly suppressed. Of note, CD9^+^Il1rn^+^ macrophages, which is transcriptionally unique relative to other clusters, were most largely found in infarcted hearts of Nlrx1KO mice, suggesting that the presence of CD9^+^Il1rn^+^ infarct macrophages is associated with loss of Nlrx1.

CD9^+^ macrophage has been suggested to exhibit a pro-fibrogenic phenotype, and expand in the early course of liver cirrhosis of both mice and human.^21^ In the setting of LPS-induced macrophage activation and lung inflammation, CD9 of macrophage negatively regulates LPS response by preventing formation of LPS receptor complex and decreasing macrophage infiltration and TNF-α production.^22^ However, CD9^+^ macrophage has not been reported in the context of infarcted heart and its role in infarct healing is largely unknown. IL-1β, as a hallmark of proinflammatory macrophage, represents a key cytokine of infarct inflammation, and the biological effects of IL-1β is counteracted by IL-1 receptor antagonist (IL-1RA), which is an endogenous inhibitor of IL-1 receptor signaling and functions as anti-inflammatory cytokine.^23^ Phase II clinical trials show promising data with anakinra, recombinant IL-1 receptor antagonist, in patients with ST-segment–elevation acute MI or heart failure with reduced ejection fraction.^24^ Here, *IL-1β* expression was significantly downregulated in CD9^+^Il1rn^+^ macrophages, whereas, *Il1rn*, the gene encoding IL-1RA, was upregulated by CD9^+^Il1rn^+^ macrophages. It has been demonstrated that *Spp1* (encoding OPN) expression was remarkably increased following infarction, and OPN is almost exclusively produced by galectin-3^hi^CD206^+^ macrophages in the infarcted heart. OPN-producing macrophages contribute to infarct repair by promoting fibrosis and clearance of apoptotic cells, and *Spp1* knockout mice are more vulnerable to develop left ventricular chamber dilatation after MI.^25^ Moreover, SPP1^+^ tumor associated macrophages showed higher reparative signatures and preferential expression of genes involved in angiogenesis.^26^ Of note, analysis of human myocardial infarction tissue specimens revealed that SPP1^+^ macrophages show increased phagocytic activity, and the presence of SPP1^+^ macrophages better predicted all fibroblasts states across different stages of human cardiac tissue remodeling.^27^ GPNMB, also known as osteoactivin, is a highly-glycosylated type I trans-membrane protein localized to the cell surface and phagosomal membranes. It has been suggested that endogenous GPNMB serves as an inflammatory stop signal inhibiting activation of T lymphocytes,^28^ and acts as a negative regulator of macrophage inflammatory responses.^29,30^ ATP6V0d2 has recently been identified as a macrophage-specific subunit of vacuolar ATPase, and inhibits macrophage inflammation and bacterial infection by promoting autophagosome and lysosome fusion,^31^ whereas Atp6v0d2-deficient macrophages exhibit augmented mitochondrial damage, enhanced inflammasome activation and compromised clearance of bacterial.^32^

Taken together, our study with single cell RNA-seq analysis identified, for the first time, a novel cluster of CD9^+^ Il1rn^+^ macrophages in the infarcted heart, in addition to subsets of Isg^+^ macrophages,^14^ CCR2^+^ macrophages,^18,19^ proliferative macrophages,^20^ and LYVE1^+^ cardiac resident macrophages.^15^ Previous studies regarding the role of Cd9, Il1rn, Gpnmb, Spp1 and Atp6v0d2 in macrophage, along with our data suggest an anti-inflammatory and reparative phenotype of CD9^+^Il1rn^+^ macrophages, which contribute to inflammation resolution, reparative fibrosis and infarct healing, accounting for preserved ventricular function and decreased mortality in Nlrx1KO mice.

### 2. The role of Nlrx1 in cardiac ischemic injury and repair

The role of Nlrx1 in immune response against pathogen infection is controversial, as experimental studies have produced conflicting results. Although originally suggested to be a negative regulator of innate immune responses to viruses, accumulating evidence shows that Nlrx1 exert diverse regulatory effects on innate immune responses depending on cell context and experimental conditions. It has been shown that Nlrx1 promotes macrophage type I IFN signaling, and both Nlrx1^−/−^ macrophages and Nlrx1-deficient mice exhibit impaired type I IFN production and enhanced viral replication.^4^ While suppressing MAVS-mediated activation of IRF3, Nlrx1 conversely facilitates virus-induced increases in IRF1 expression via opposing regulatory mechanisms and thereby overally promotes early innate immune antiviral defense in hepatocytes.^5^ Also, rather than inhibiting NF-κB signaling, Nlrx1 has been shown to potentiate the activation of NF-κB and JNK in response to several stimuli,^6^ although it is originally suggested to be a potent inhibitor of NF-κB activation and IFNβ production in response to sendai virus and sindbis virus.^3^ It has been demonstrated that Nlrx1 is hijacked by *Listeria* to evade killing both in vitro and in vivo, and *Listeria*-induced mortality was significantly lower in Nlrx1^-/-^ mice.^7^ Recent studies suggest that Nlrx1 positively correlates with simian immune-deficiency virus viremia in nonhuman primates^33^ and with HIV-1 viremia in patients,^34^ and multi-omics analyses reveal that Nlrx1 enhances oxidative phosphorylation and glycolysis during HIV-1-infection of CD4^+^ T cells to promote viral replication.^34^

Again, current available studies regarding the involvement of Nlrx1 in tissue injury and repair also yield controversial results. The studies by Kors’s group suggested that Nlrx1 dampens oxidative stress and apoptosis via control of mitochondrial activity, and loss of Nlrx1 results in increased oxidative stress and apoptosis in epithelial cells during renal ischemia-reperfusion injury.^8^ However, another study by the same group showed that deletion of Nlrx1 increases fatty acid metabolism and prevents diet-induced hepatic steatosis and metabolic syndrome.^9^ It has been shown that Nlrx1 expression was decreased in patients with chronic obstructive pulmonary disease, and Nlrx1 suppression correlated directly with disease severity. In murine models, cigarette smoke inhibited Nlrx1, and cigarette smoke-induced alveolar destruction and inflammasome activation were augmented in Nlrx1-deficient animals.^35^ Overall, these results suggest that the role of Nlrx1 in tissue injury and repair depends on tissue context and the pathogenetic condition or disease model used.

It has been suggested that Nlrx1 is highly expressed in the heart tissues,^3^ and consistently, mining of The Human Protein Atlas revealed high expression of Nlrx1 in heart and cardiomyocytes. Our study confirmed that protein expression of Nlrx1 is remarkably high in primary cardiomyocytes and H9C2 myocyte cell line, followed by RAW264.7 macrophages cell line, and lowest in bone marrow derived macrophages. Moreover, Nlrx1expression is significantly increased in the infarcted heart in vivo, however, the specific role and functional significance of Nlrx1 in myocardial ischemic injury and healing is poorly understood. It has been shown that Nlrx1 deletion increases ischemia-reperfusion damage and activates glucose metabolism in the isolated mouse heart.^36^ In contrast to this observation, our in vivo results show that loss of Nlrx1 attenuates ventricular dysfunction and decreases post-infarction mortality, which is associated with increased CD9^+^IL1rn^+^ macrophages in the Nlrx1KO infarcted hearts, and CD9^+^CD11b^+^ macrophages exhibit a reparative and anti-inflammatory phenotype with upregulation of wound healing associated genes and downregulation of proinflammatory genes. Given the high expression of Nlrx1 in cardiomyocytes relative to that in macrophages, the presence of CD9^+^IL1rn^+^ reparative macrophages may be largely attributed to deficiency of Nlrx1 in cardiomyocytes rather than macrophage itself, although the potential effects of Nlrx1 loss in other cell types can’t be excluded. This leads to the hypothesis that certain mediator(s) secreted by Nlrx1 deficient cardiomyocytes upon ischemic injury, may modulate and reprogram infarct macrophages towards a reparative and/or anti-inflammatory phenotype, thereby contributing to infarct healing, preserved cardiac function and increased post-infarction survival in Nlrx1 deficient mice.

### 3. Stressed cardiomyocyte-derived mediators regulate phenotype and function of macrophages in the infarcted heart

Although the effects of infiltrating macrophages on survival of cardiomyocytes and healing of injured hearts have been intensely studied,^1,2^ whether or not and how stressed cardiomyocytes might influence the phenotype and function of macrophages in the infarcted heart was largely unexplored. In the setting of MI, cardiomyocytes have long been viewed as passive targets of ischemic injury, and dying cardiomyocytes release danger signals triggering an innate immune response.^37,38^ It is also worth noting that bordering areas of the infarcted myocardium can also experience impaired blood supply and reduced oxygen delivery, leading to altered metabolic and mechanical processes, and in turn the living cardiomyocytes under stress along infarct border zone may actively secrete mediators which could selectively recruit specific subsets of monocytes, or guide macrophages activation towards anti-inflammatory and/or reparative phenotype to restrain local inflammation and boost infarct healing.^37^ This notion has been proofed by an excellent study published in Nature Medicine, Lörchner and co-workers demonstrated that Reg3b, a small secretory protein released by cardiomyocytes of infarct border zone, exert a crucial role in post-infarction inflammation resolution and infarct repair by recruiting reparative macrophages that promote matrix preservation.^39^ Additionally, cross-talk between cardiomyocytes and fibroblast has also gained attention recently, and PCSK6 (proprotein convertase subtilisin/kexin type 6) has been identified as a novel cardiomyocytes-derived mediator secreted by hypoxic cardiomyocytes and elevated in infarcted hearts, potentiating collagen production by fibroblasts in the infarcted hearts.^40^

In our study, conditioned medium from Nlrx1 deficient cardiomyocytes upon hypoxia skewed macrophage transition towards a reparative and inflammation-suppressed phenotype, and gain-and loss-of Nlrx1 study combined with unbiased RNA-seq analysis revealed that CCN2 was significantly upregulated in NLRX1 knockdown cardiomyocytes, but remarkably downregulated in NLRX1-overexpressing cardiomyocytes in response to hypoxia stimulation. Both protein expression of CCN2 and secreted CCN2 in the supernatant were increased in Nlrx1 knockdown cardiomyocytes upon hypoxia stimulation. Moreover, CCN2 protein was significantly elevated in the infarcted hearts of Nlrx1KO mice in vivo. Furthermore, exogenous CCN2 partially mediates macrophage polarization towards a reparative phenotype. Taken together, CCN2, which is significantly elevated in the Nlrx1KO infarcted hearts and secreted by Nlrx1 deficient cardiomyocytes upon hypoxia, is identified as a cardiomyocytes-derived mediator responsible, at least partly, for reprograming macrophages towards a reparative and anti-inflammatory phenotype contributing to infarct healing.

Moreover, consistent with our present observations, previous studies have shown that cardiac CCN2 expression is elevated following myocardial ischemia^41^ and exert cardioprotective role in the setting of MI,^42^ ischemia-reperfusion injury,^43^ and chronic pressure overload^44^ through undetermined mechanisms. Our study adds to the understanding of biological function of CCN2 and provides novel insights into cardioprotective effect of CCN2 by showing that cardiomyocytes-derived CCN2 can modulate transition of infarct macrophages towards a reparative phenotype. However, considering the repertoire of mediators secreted by cardiomyocytes in the infarct border zone, an exclusive role for CCN2 in modulating phenotype and function of infarct macrophages is unlikely. Other cardiomyocytes- and non-cardiomyocytes-derived mediators, yet to be identified, may orchestrate the phenotype and function of infarct macrophages driving reparative, fibrogenic, or angiogenic responses.

In conclusion, our data together with published results demonstrating that cardiomyocytes-secreted Reg3b facilitates inflammation resolution and infarct repair by recruiting reparative macrophages, establish a new paradigm for the cross-talk between cardiomyocytes-derived mediators and phenotypic plasticity of cardiac macrophages in the infarcted hearts. These results also reinforce the concept that instead of passive targets of ischemic injury, the living cardiomyocytes along infarct border-zone may actively secrete mediators which potentially orchestrate infiltrating dynamics and functional phenotype of infarct macrophages driving reparative responses to heal the infarcted hearts.

## Acknowledgements

This work was supported by the National Natural Science Foundation of China (82170249, 82070051, 81650022, 81602494, 81772167, 81971563 and 82272330).

---

**Supplementary Figure S1:**
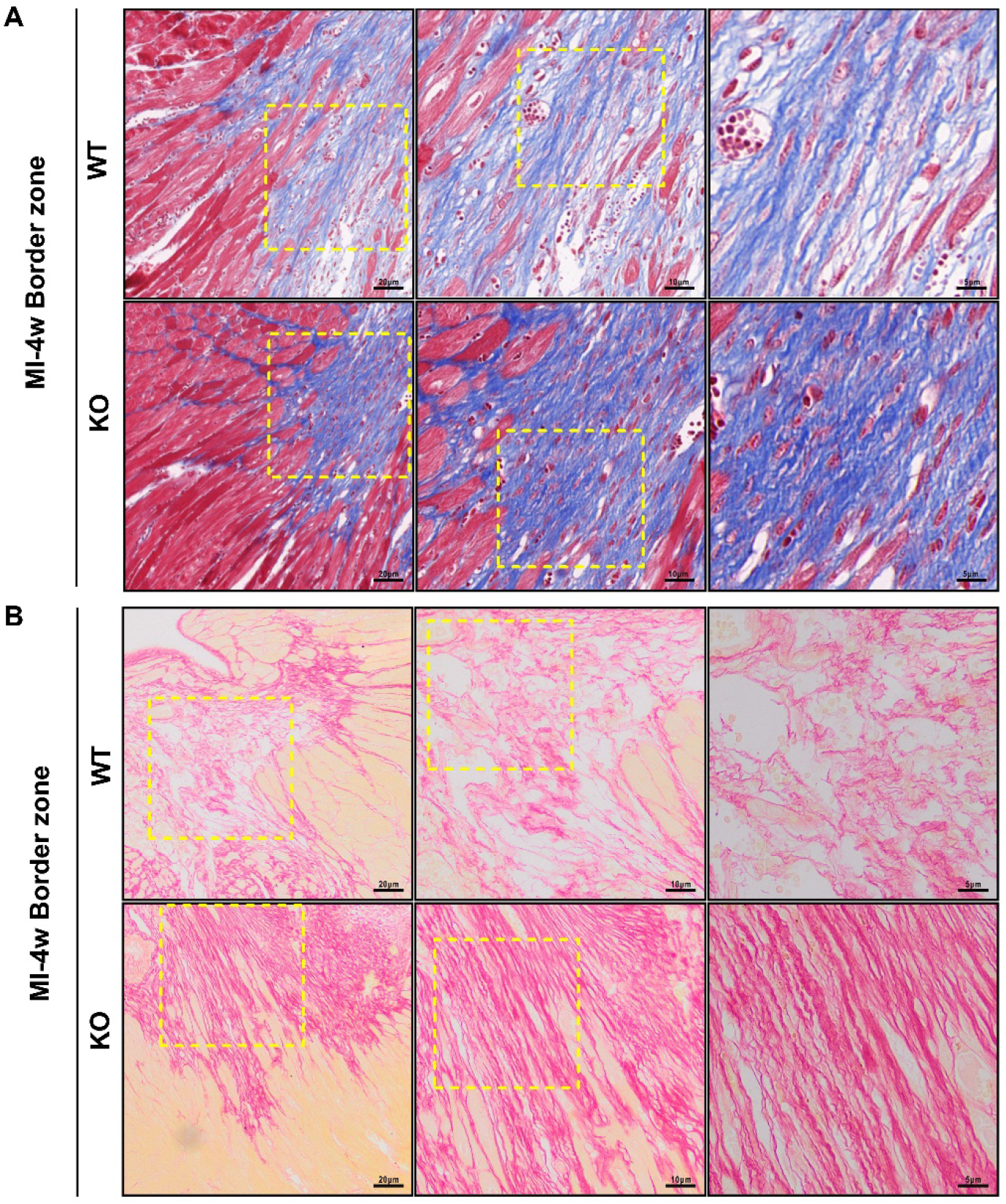
Representative images of masson’s trichrome (**A**) and picrosirius red staining (**B**) showing collagen deposition in the infarct border zone 4 weeks after MI. Nlrx1KO infarct border zone was characterized by closely assembled and well-aligned collagen fibers, in contrast to loosely and fragmented collagen fibers in the infarct border zone of WT mice. Scale bar as indicated.

**Supplementary Figure S2:**
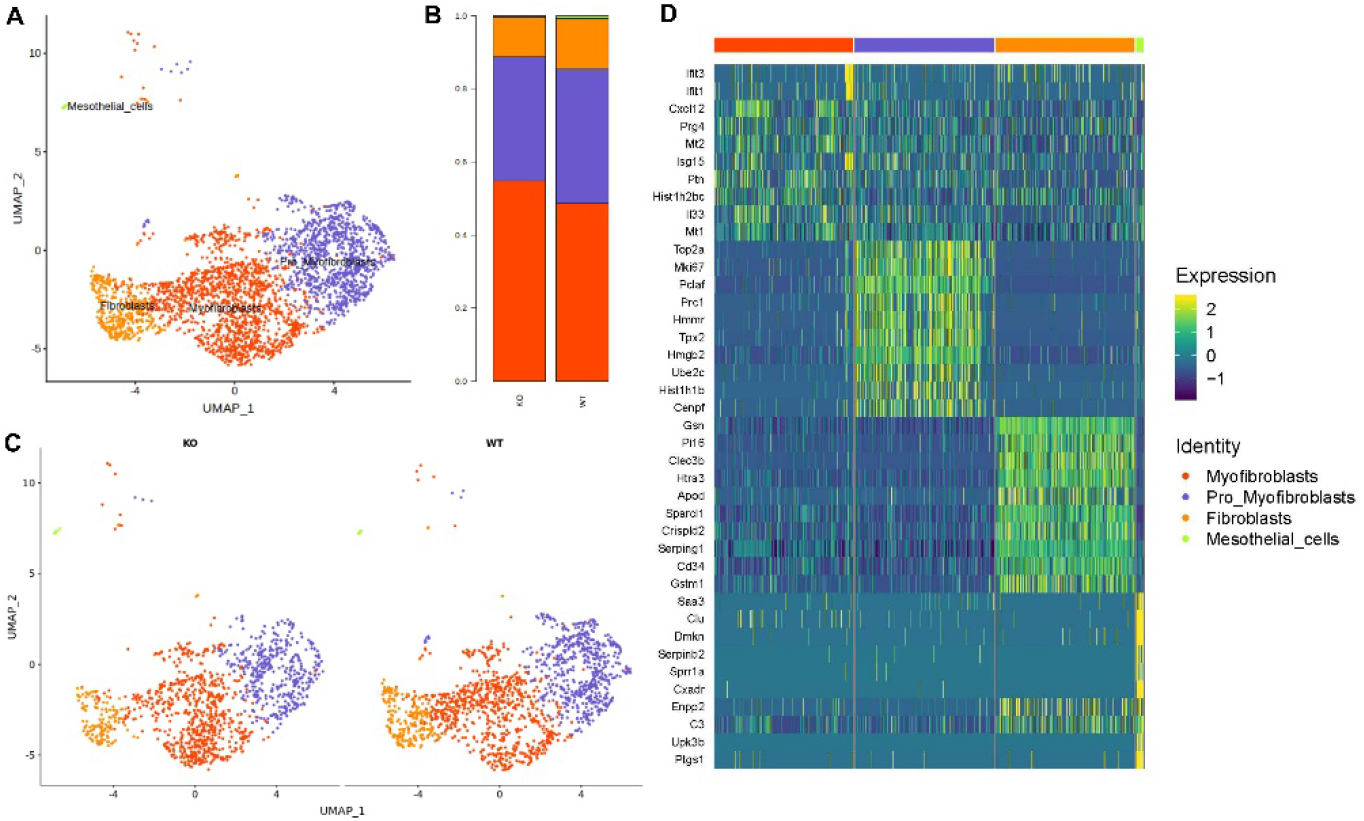
Single-cell RNA-seq analysis of fibroblasts in the infarcted hearts. **A**, UMAP plot depicting 4 sub-clusters of fibroblasts in infarcted hearts 3 days after infarction. **B**, Fraction of each subset of fibroblasts in WT and Nlrx1KO group. Fibroblasts, myofibroblasts, pro-myofibroblasts and mesothelial cells, accounted for 48.6-54.8%, 34-36.8%, 10.8-13.9%, and 0.4-0.7% of all sequenced fibroblasts respectively. **C**, UMAP plot depicting sub-clusters of fibroblasts annotated by group. **D**, Heatmap of the top 10 differentially expressed genes in each subset of fibroblasts (top, color-coded by subset).

**Supplementary Figure S3:**
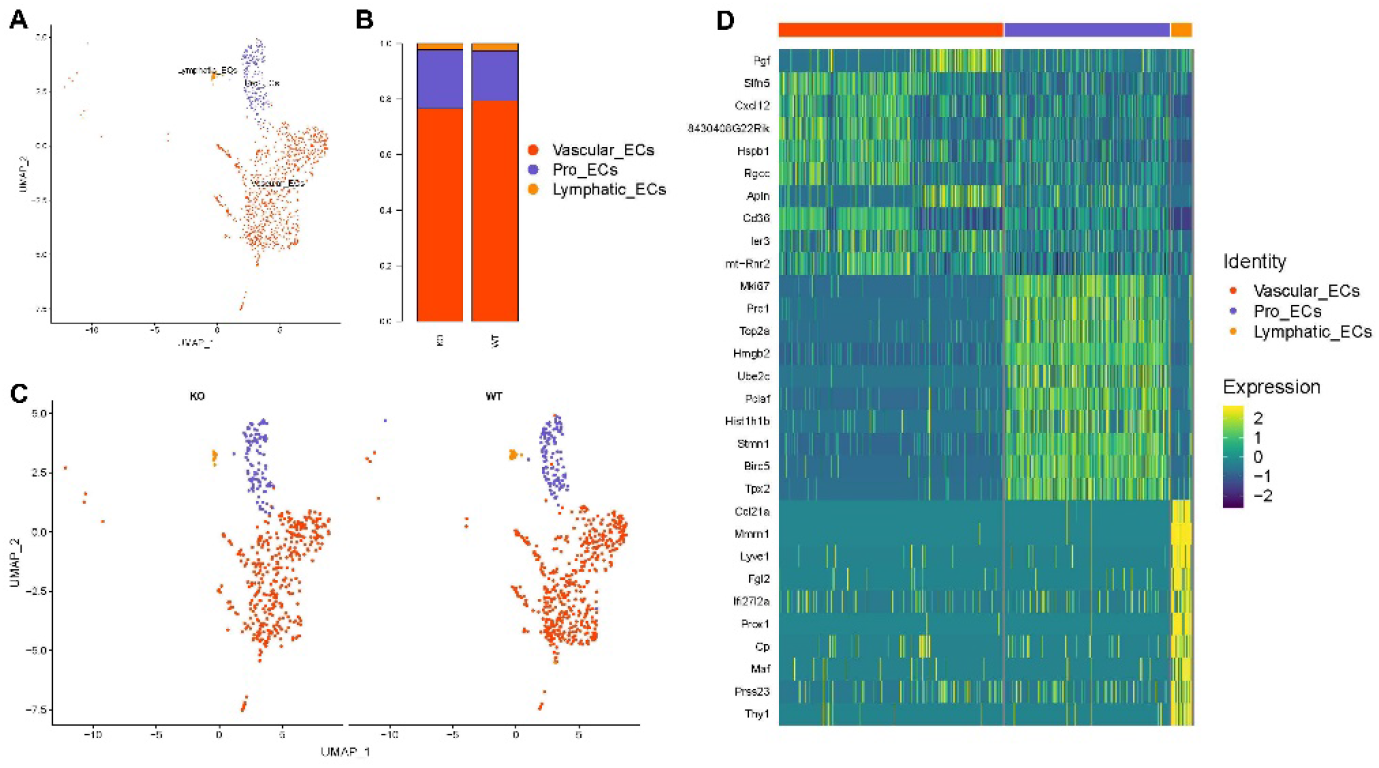
Single-cell RNA-seq analysis of endothelial cells (ECs) in the infarcted hearts. **A**, UMAP plot depicting 3 sub-clusters of ECs of infarcted hearts 3 days after infarction. **B**, Fraction of each subset of ECs in WT and Nlrx1KO group. Vascular ECs, proliferative ECs and lymphatic ECs accounted for 76.6-79.4%, 18-21.1%, and 2.4% of all sequenced ECs respectively. **C**, UMAP plot depicting sub-clusters of ECs annotated by group. **D**, Heatmap of top 10 differentially expressed genes in each subset of ECs (top, color-coded by subset).

**Supplementary Figure S4:**
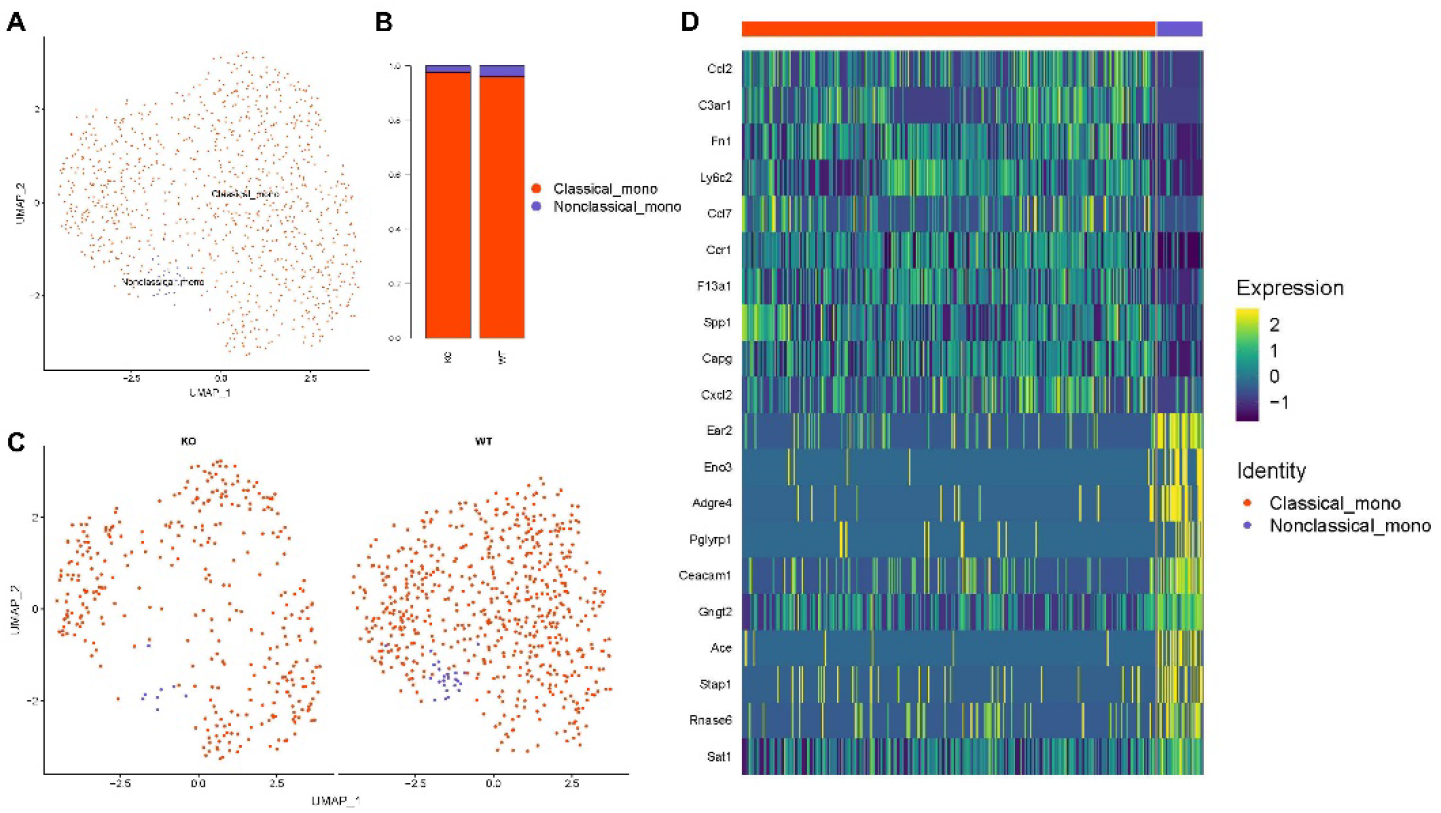
Single-cell RNA-seq analysis of monocytes in the infarcted hearts. **A**, UMAP plot depicting 2 sub-clusters of monocytes of infarcted hearts 3 days after infarction. **B**, Fraction of classical and nonclassical monocytes in WT and Nlrx1KO group. **C**, UMAP plot depicting sub-clusters of monocytes annotated by group. **D**, Heatmap showing top 10 differentially expressed genes in each subset of monocytes.

**Supplementary Figure S5:**
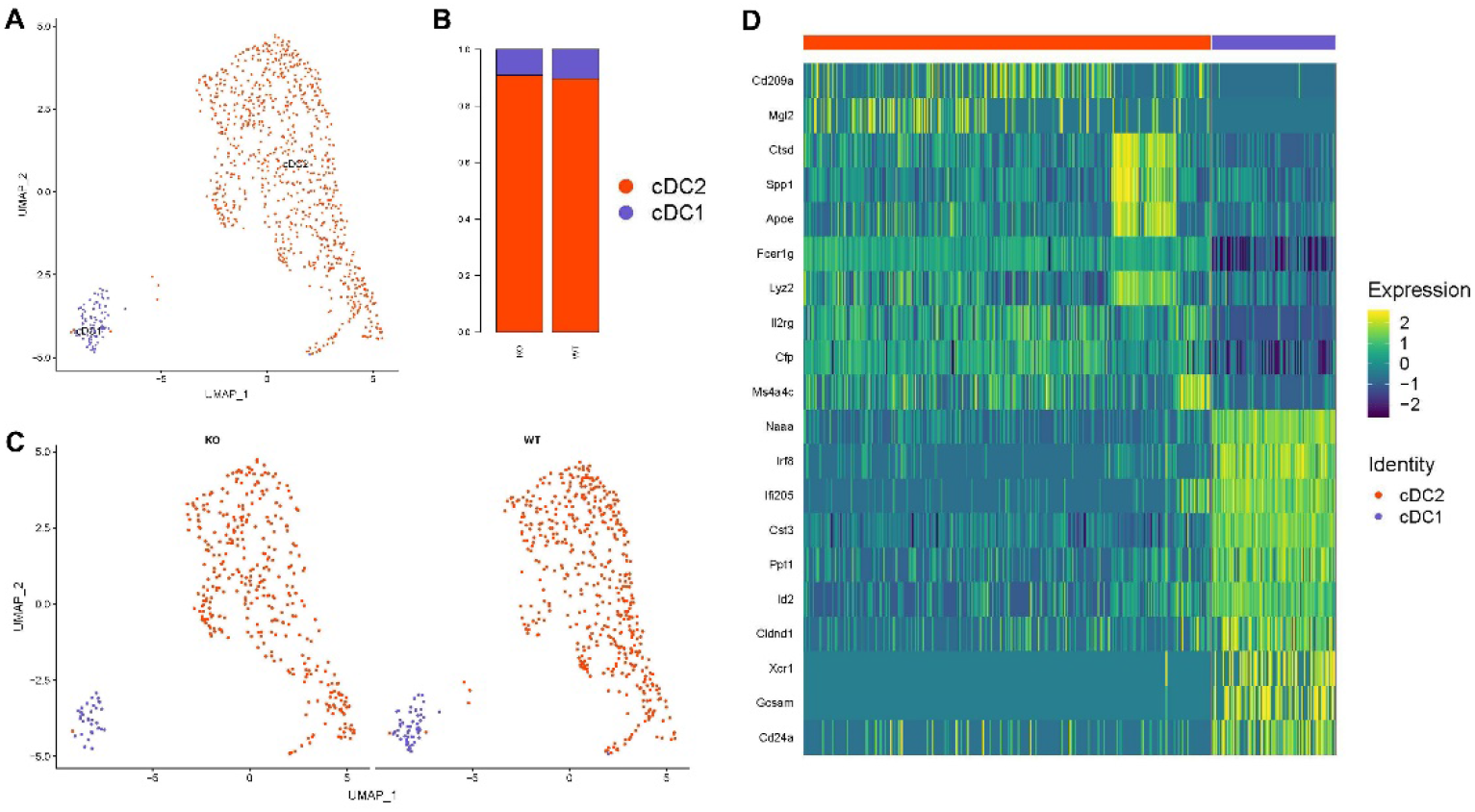
Single-cell RNA-seq analysis of DCs in the infarcted hearts. **A**, UMAP plot depicting 2 sub-clusters of conventional dendritic cells (DCs) of infarcted hearts 3 days after infarction. **B**, cDC1 and cDC2 accounted for 10% and 90% of DCs, respectively. **C**, UMAP plot depicting sub-clusters of DCs annotated by group. **D**, Heatmap of top 10 differentially expressed genes in each subset of DCs.

**Supplementary Figure S6:**
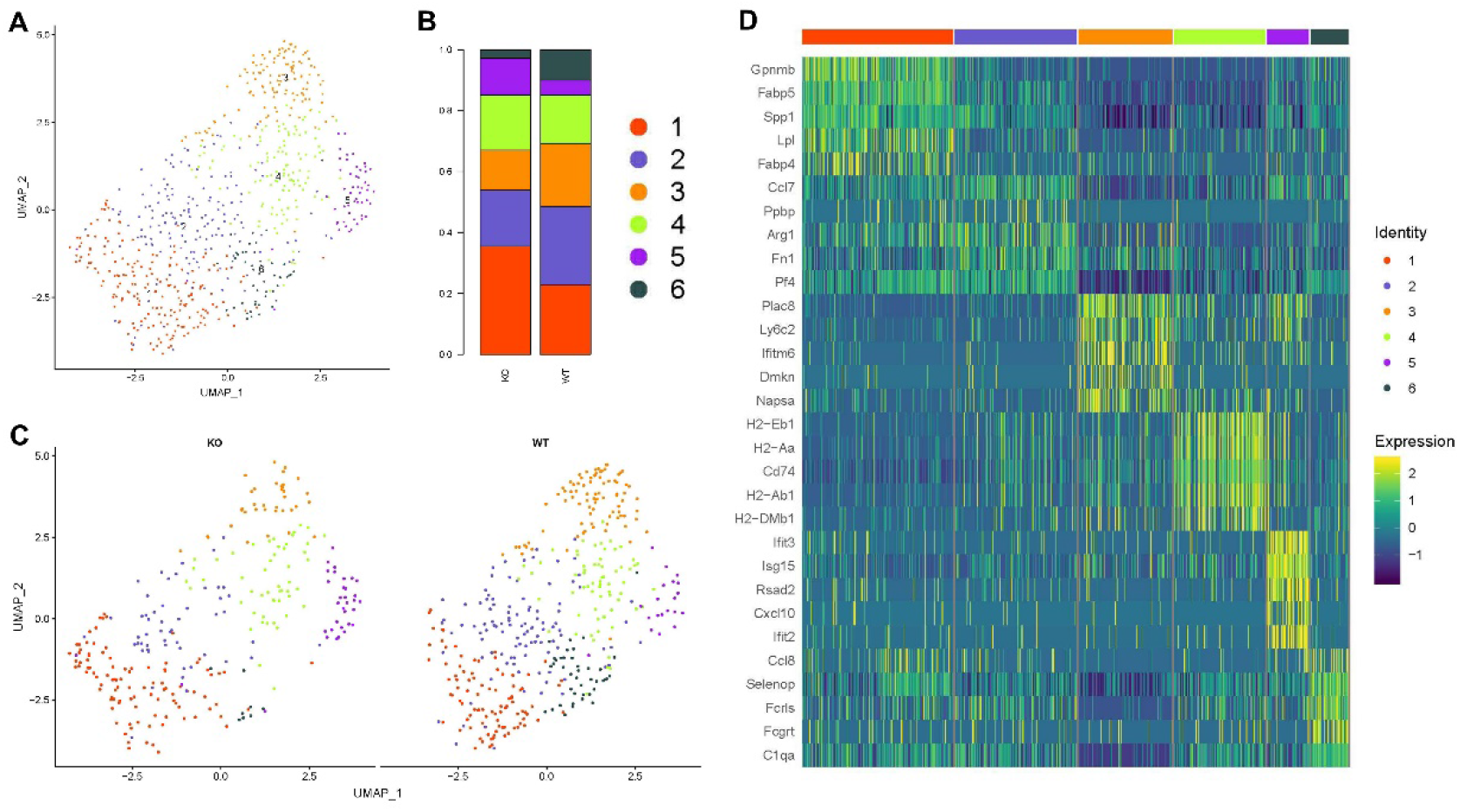
Single-cell RNA-seq analysis of neutrophils in the infarcted hearts. **A**, UMAP plot depicting 6 sub-clusters of neutrophils of infarcted hearts 3 days after infarction. **B**, Fraction of each subset of neutrophils in WT and Nlrx1KO group. **C**, UMAP plot depicting sub-clusters of neutrophils annotated by group. **D**, Heatmap showing top 5 differentially expressed genes in each subset of neutrophils (top, color-coded by subset).

**Supplementary Figure S7:**
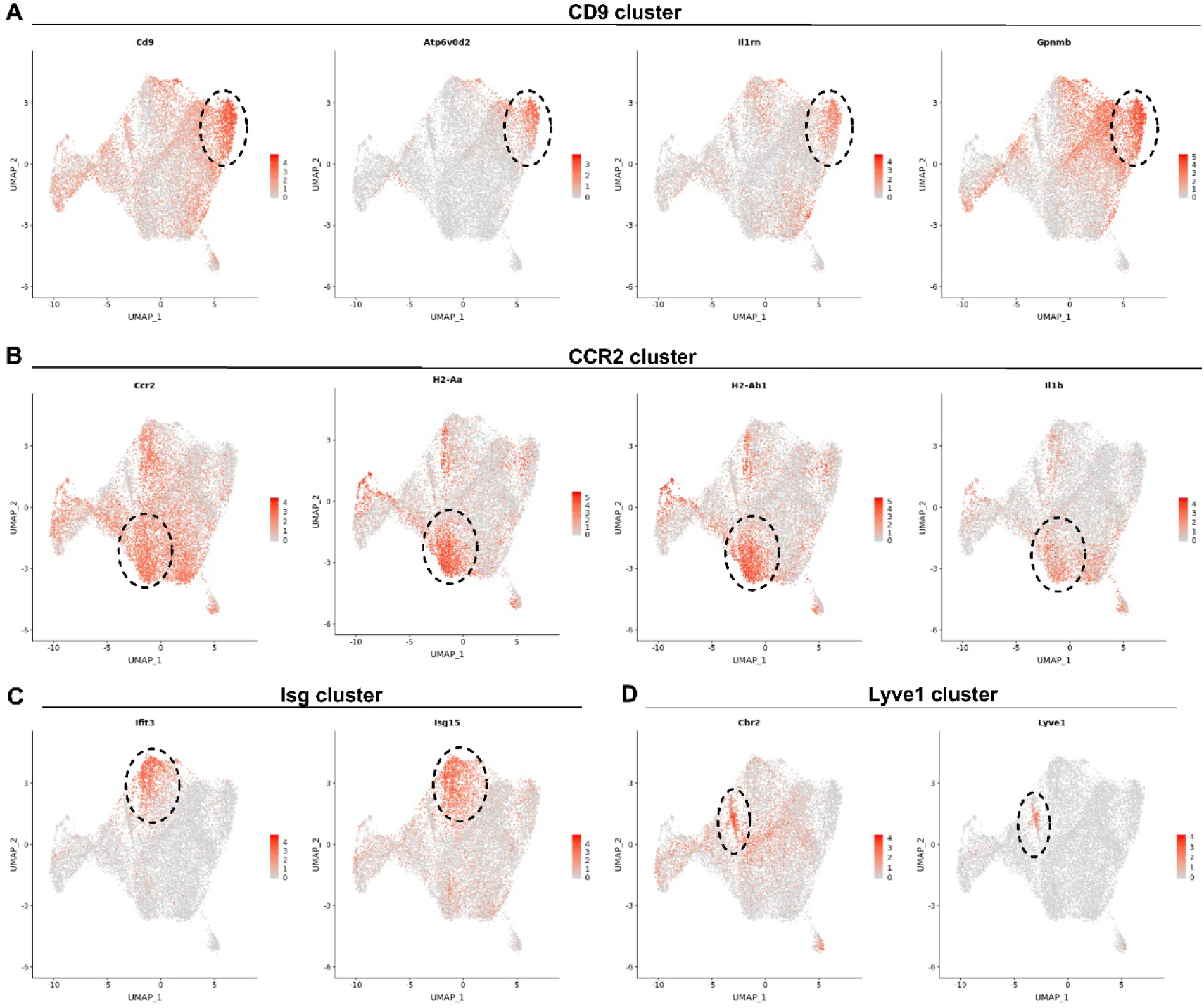
Feature plots depicting single-cell gene expression of cluster-defining signatures of macrophage subsets. **A**, CD9 cluster of macrophages showed high expression of wound healing and anti-inflammation associated genes including *CD9, Atp06vd2, IL1rn* and *Gpnmb*. **B**, CCR2 cluster of macrophages were characterized by upregulation of *CCR2, IL1b, H2-Aa* and *H2-Ab1*. **C**, Isg cluster of macrophages exhibited increased expression of *ifit3* and *Isg15*. **D**, Lyve1 cluster of cardiac resident macrophage showed exclusively high expression of *Lyve1* and relatively increased *Cbr2*.

**Supplementary Figure S8:**
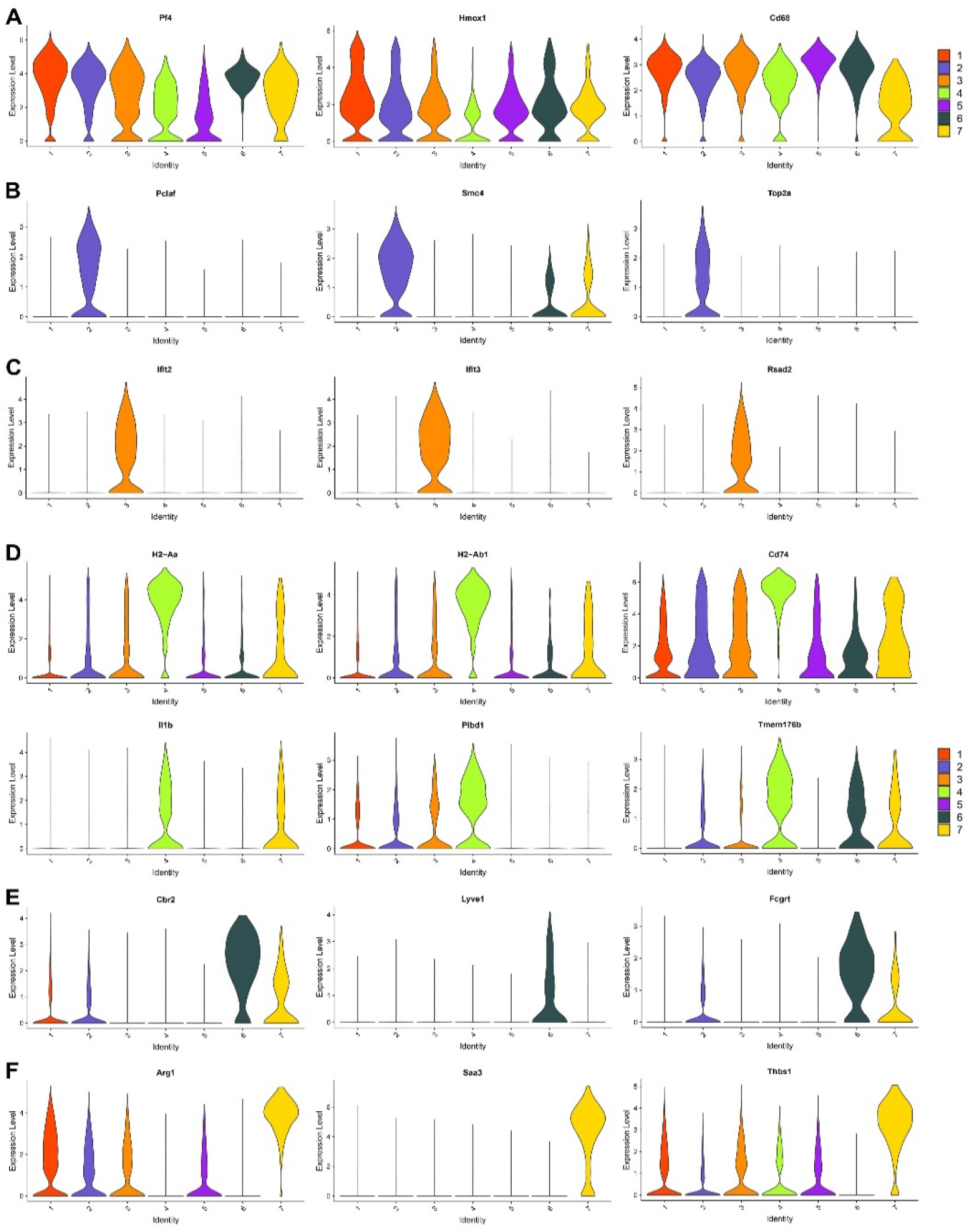
Violin plots depicting expression level of cluster-defining marker genes across subsets of infarct macrophage. **A**, Classic marker genes of macrophage including *CD68, Pf4* and *Hmox1* were widely expressed throughout all macrophage clusters with somewhat variability of expression level among subsets. **B**, Cluster 2 of proliferative macrophages were characterized by high expression of proliferation indicative genes like *Pclaf, Smc4*, and *Top2a*. **C**, Isg cluster exhibited significantly increased expression of interferon-stimulated genes such as *ifit2, ifit3* and *Rasd2*. **D**, CCR2 cluster were featured by high expression of major histocompatibility complex (MHC)-II associated genes (such as *H2-Eb1, H2-Aa1, H2-Ab1*), and typical proinflammatory genes like *Ccr2*, *Il1b*, *Plbd1* and *Tmem176b*. **E**, Lyve1 cluster of cardiac resident macrophage was characterized by high expression of *Lyve1, Cbr2* and *Fcgrt*. **F**, Cluster 7 of macrophages showed high expression of *Saa3, Arg1,* and *Thbs1*.

**Supplementary Figure S9:**
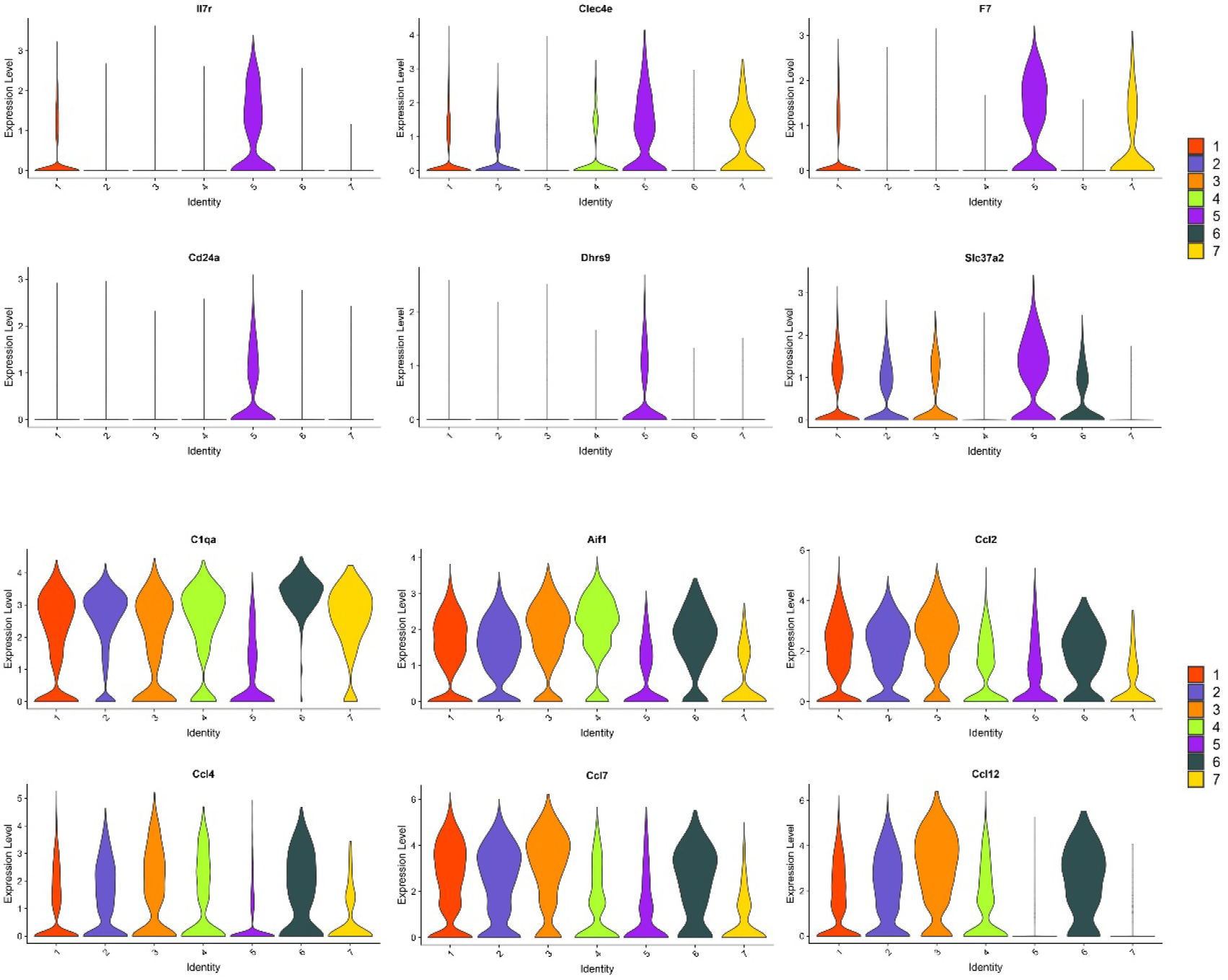
Violin plots depicting differential expression of genes across subsets of infarct macrophages. In addition to *Cd9, Il1rn, Spp1, Gpnmb* and *Atp6v0d2,* other healing and inflammation resolution associated genes including *Il7r, Clec4e, F7, CD24a, Dhrs9* and *Slc37a2* were relatively upregulated in CD9 cluster, whereas proinflammatory genes like *C1qa, Aif1,* and cytokines including *Ccl2, Ccl4, Ccl7* and *Ccl12* were downregulated in CD9 cluster relative to other clusters of infarct macrophages.

**Supplementary Figure S10:**
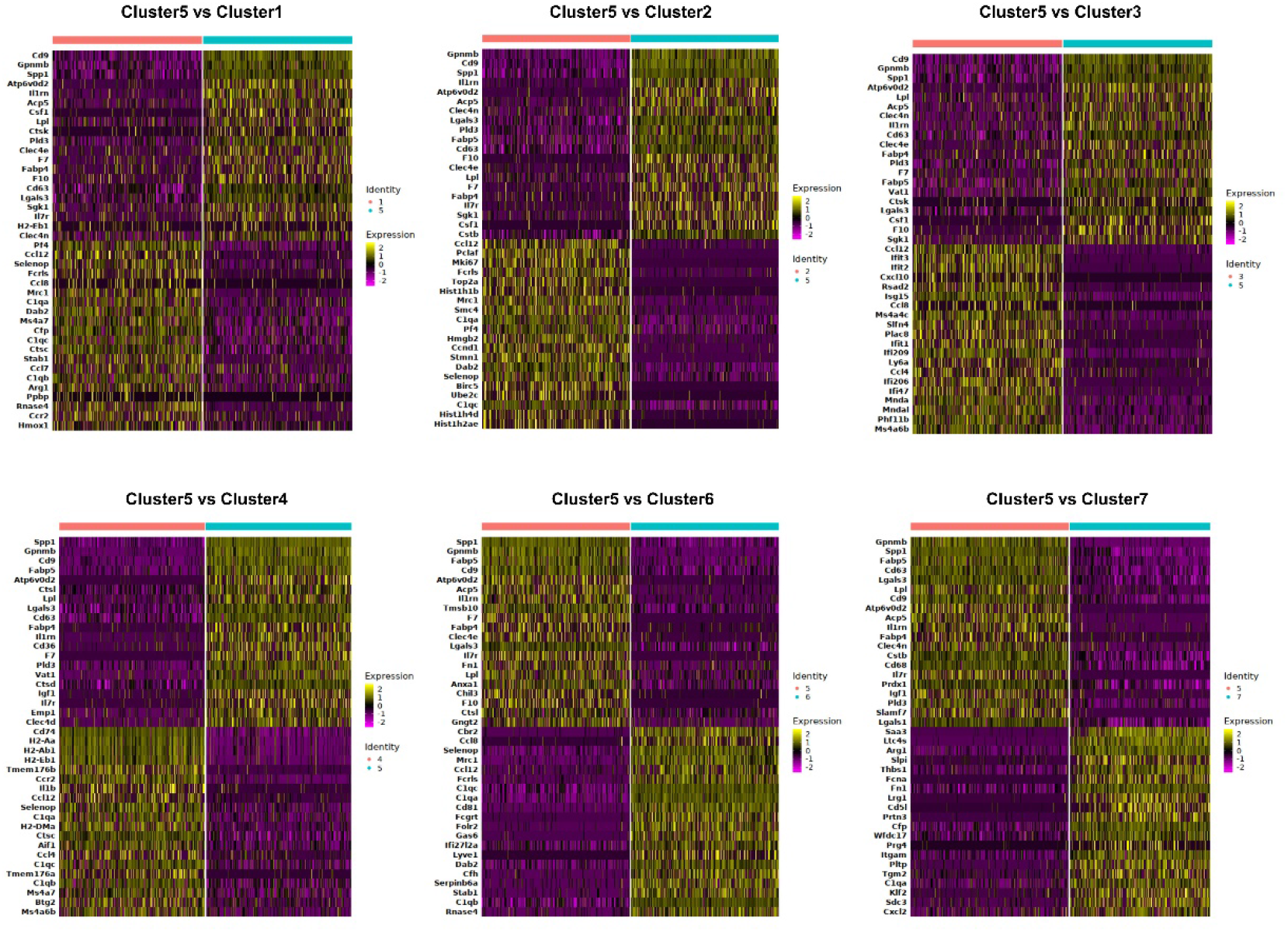
Heatmap of differentially expressed genes in CD9 cluster versus other clusters of infarct macrophages.

**Supplementary Figure S11:**
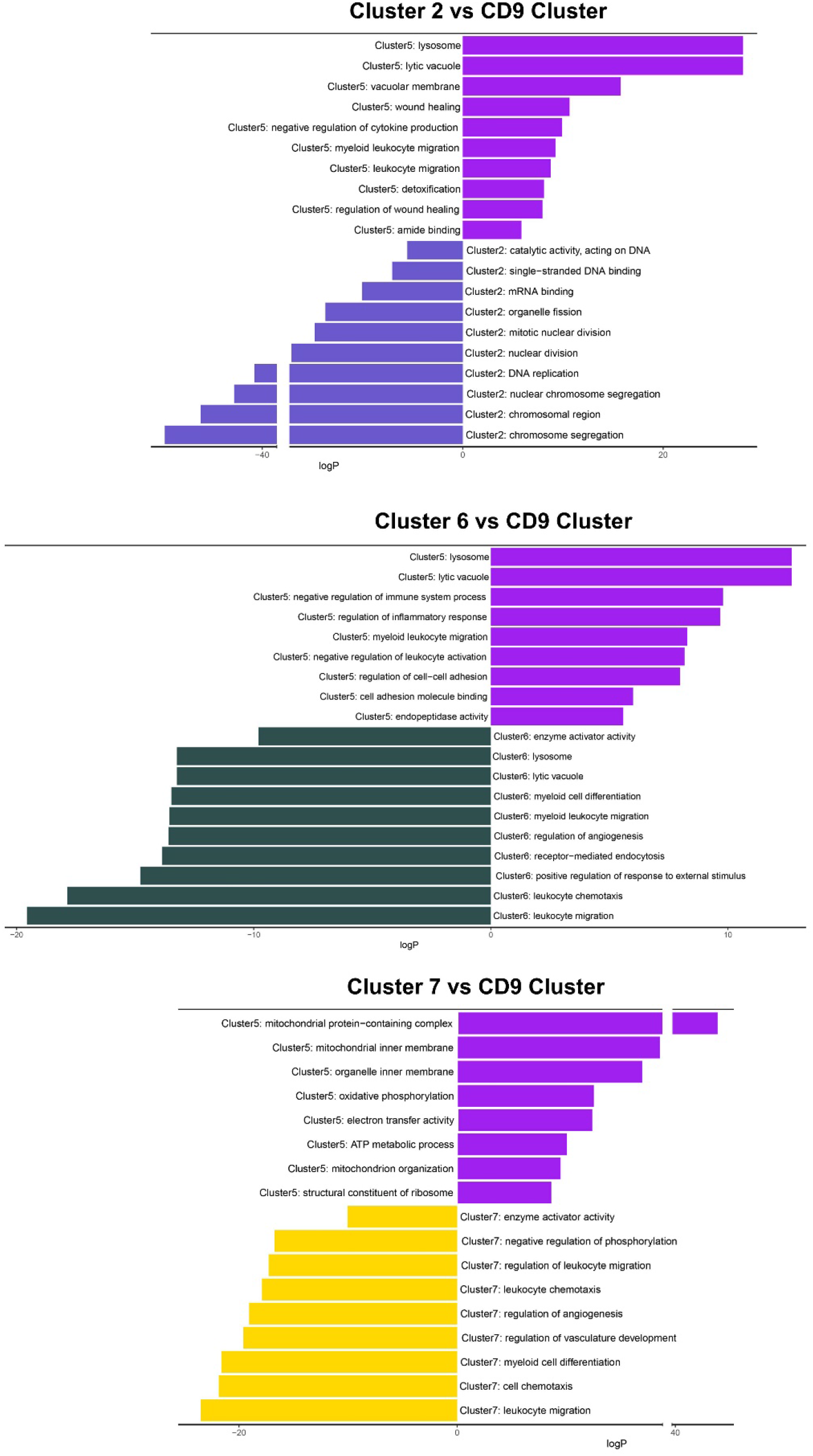
Pathway enrichment analysis of differentially expressed genes in CD9 cluster versus other clusters of macrophages. Pathway enrichment is expressed as the –log[P] adjusted for multiple comparison.

**Supplementary Figure S12:**
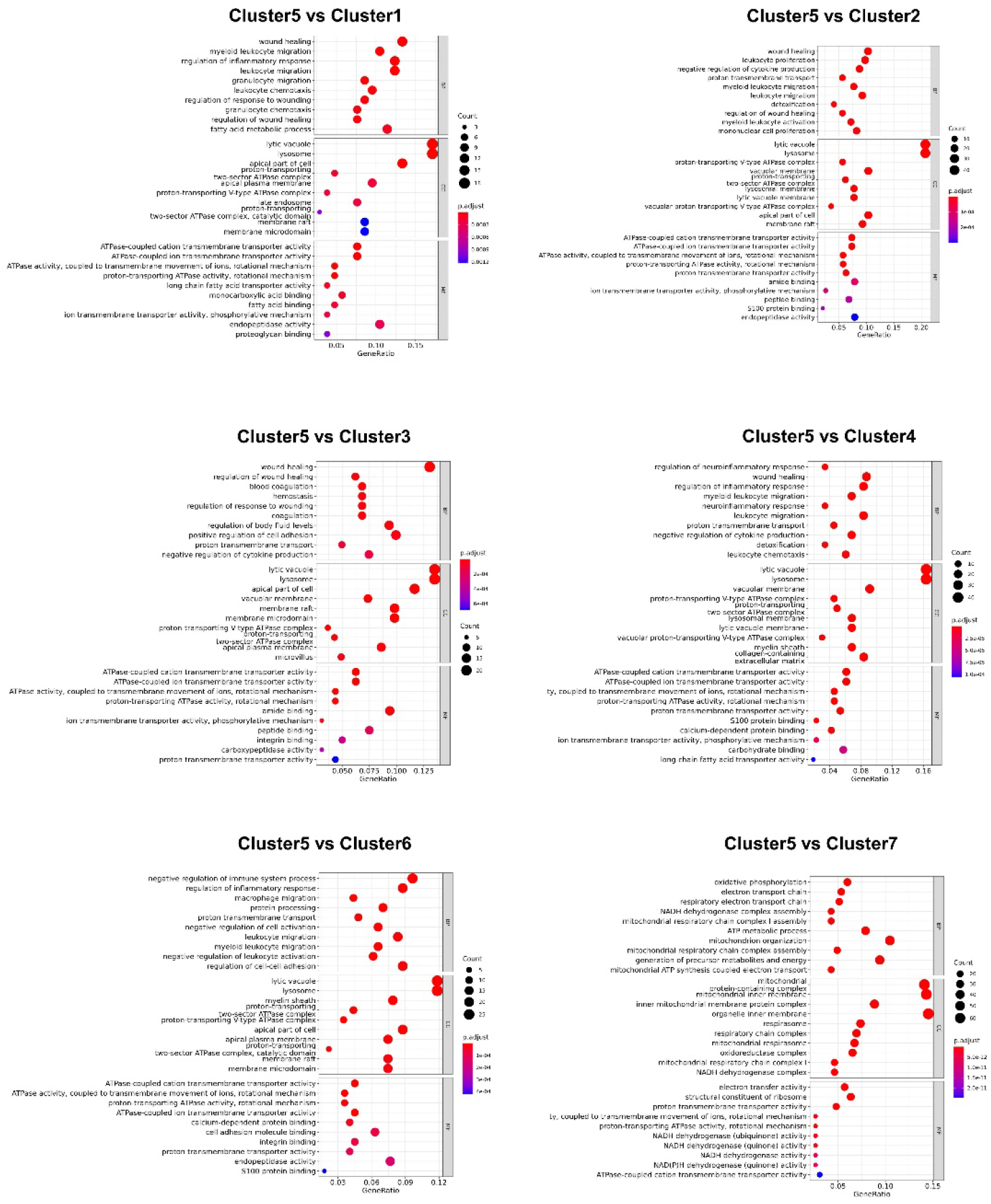
gene ontology (GO) enrichment analysis of differentially expressed genes in CD9 cluster versus other clusters of macrophages. CD9 cluster was enriched in homeostatic and reparative functions including wound healing, endocytosis, lysosome function, regulation of wound healing, negative regulation of cytokine production and inflammatory response, and detoxification relative to other clusters of infarct macrophages.

**Supplementary Figure S13:**
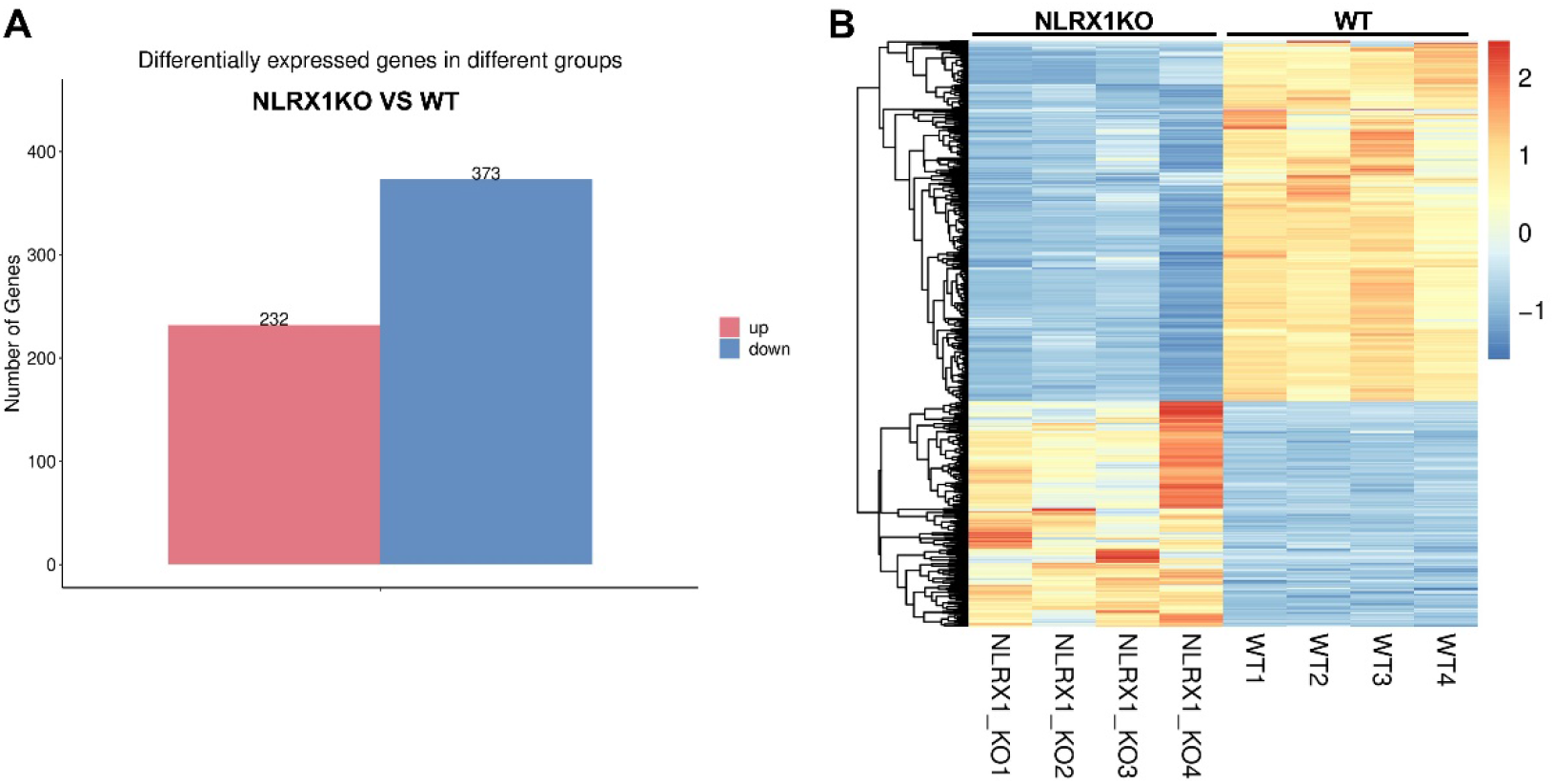
Transcriptomic comparison of CD9+CD11b+ infarct macrophages between Nlrx1KO and WT mice. **A**. Totally 232 genes were upregulated, and 373 genes were downregulated in CD9^+^CD11b^+^ macrophages of Nlrx1KO infarcts, in comparison to WT macrophages (>2-fold, padj<0.05). **B**, Heatmap showing differentially expressed genes in RNA-seq transcriptomic comparisons of CD9^+^CD11b^+^ macrophages between Nlrx1KO and WT mice.

**Supplementary Figure S14:**
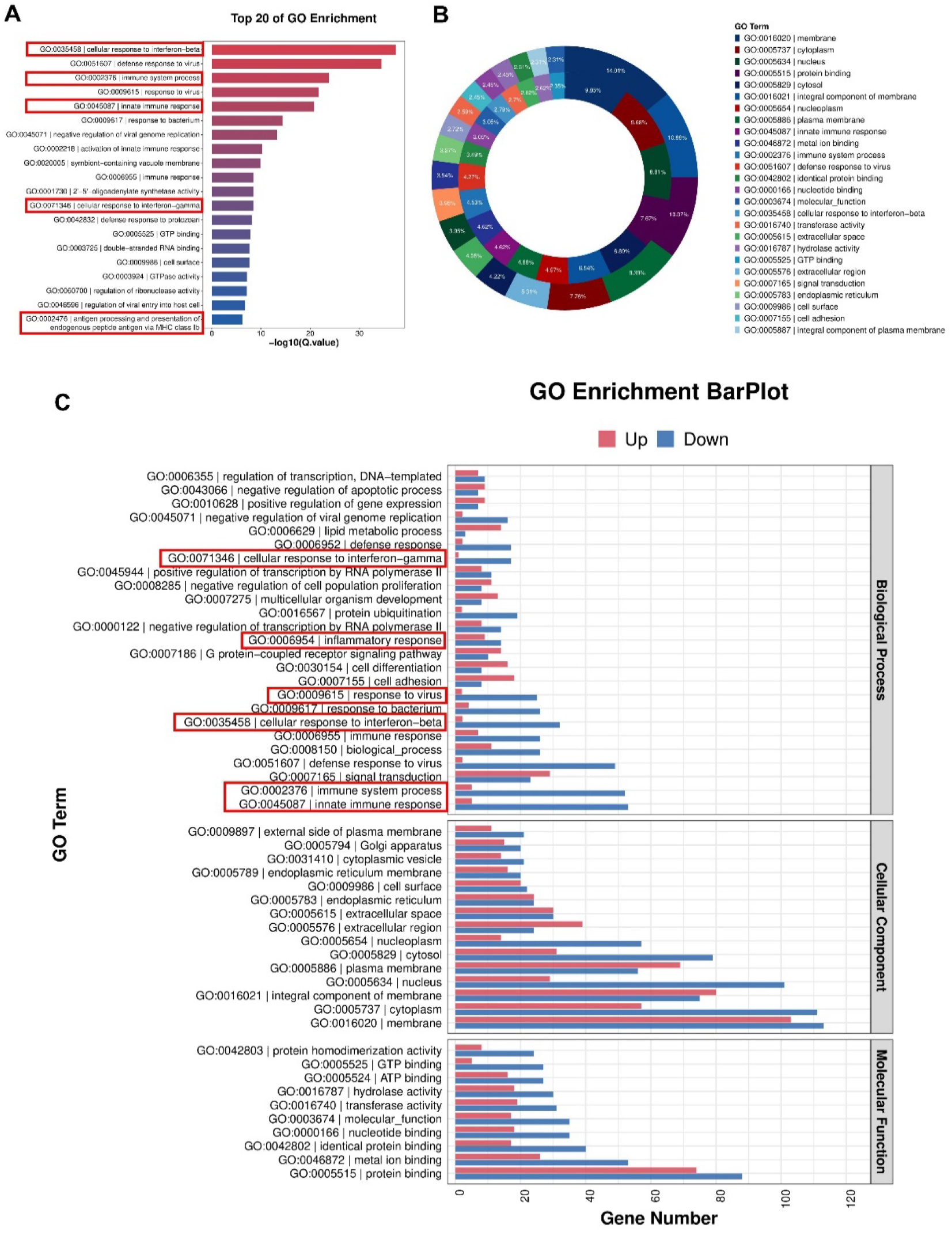
Gene ontology (GO) enrichment analysis of the identified DEGs between Nlrx1KO and WT CD9+CD11b+ macrophages. **A**, Top 20 of gene ontology (GO) enrichment analysis of the identified DEGs between Nlrx1KO and WT CD9^+^CD11b^+^ macrophages. **B**, Doughnut plot of GO enrichment analysis of the identified DEGs. **C**, Barplot showing the number of upregulated (red) and downregulated (blue) genes of each GO enrichment of biological process, cellular component, and molecular function. DEGs indicates differentially expressed genes.

**Supplementary Figure S15:**
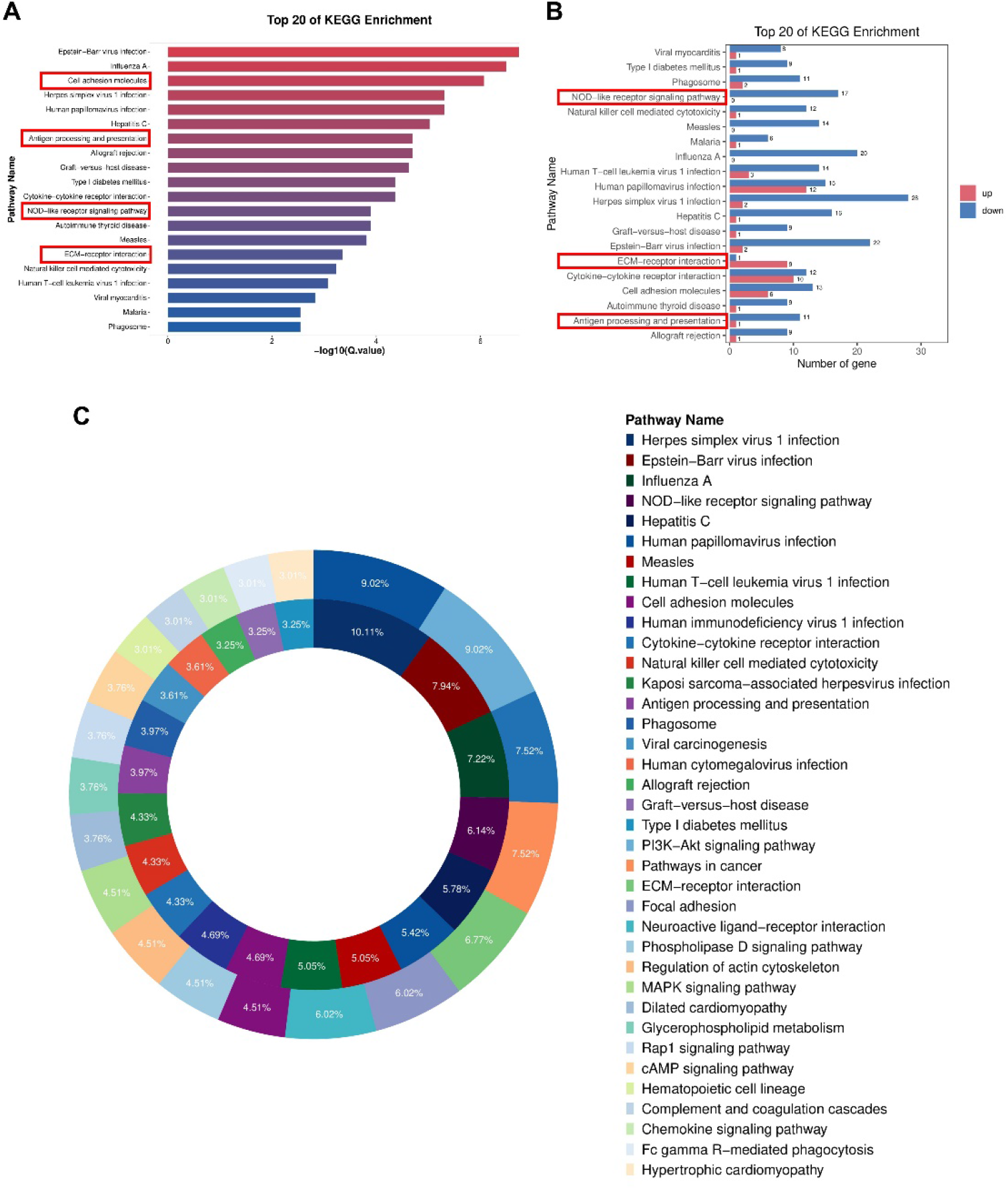
KEGG pathway enrichment analysis of the identified DEGs between Nlrx1KO and WT CD9+CD11b+ macrophages. **A**, Top 20 of KEGG pathway enrichment analysis of the identified DEGs between Nlrx1KO and WT CD9^+^CD11b^+^ macrophages. **B**, Barplot showing the number of upregulated (red) and downregulated (blue) genes of top 20 KEGG pathway. **C**, Doughnut plot of KEGG pathway enrichment analysis of the identified DEGs. DEGs indicates differentially expressed genes.

**Supplementary Figure S16:**
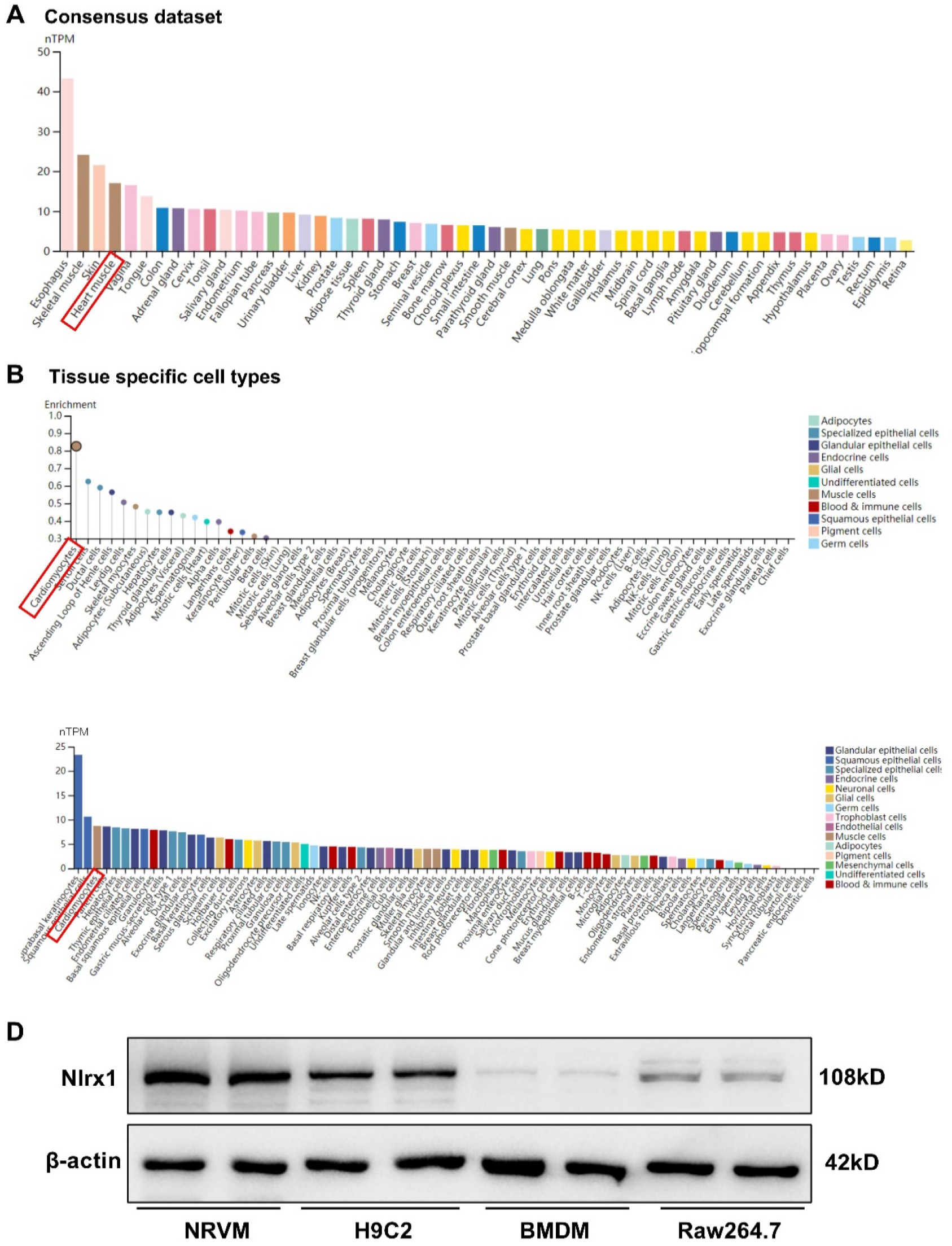
**A-C,** Human Nlrx1 is highly expressed in hearts and cardiomyocytes. Cell and tissue expression data mined from The Human Protein Atlas (www.proteinatlas.org) using Nlrx1 as the key word identifier. **D,** Protein expression of Nlrx1 in cardiomyocytes and macrophages was determined by western blotting. Nlrx1 expression was remarkably high in primary cardiomyocytes and H9C2 myocyte cell line, followed by RAW264.7 macrophages cell line, and lowest in bone marrow derived macrophages.

**Supplementary Figure S17:**
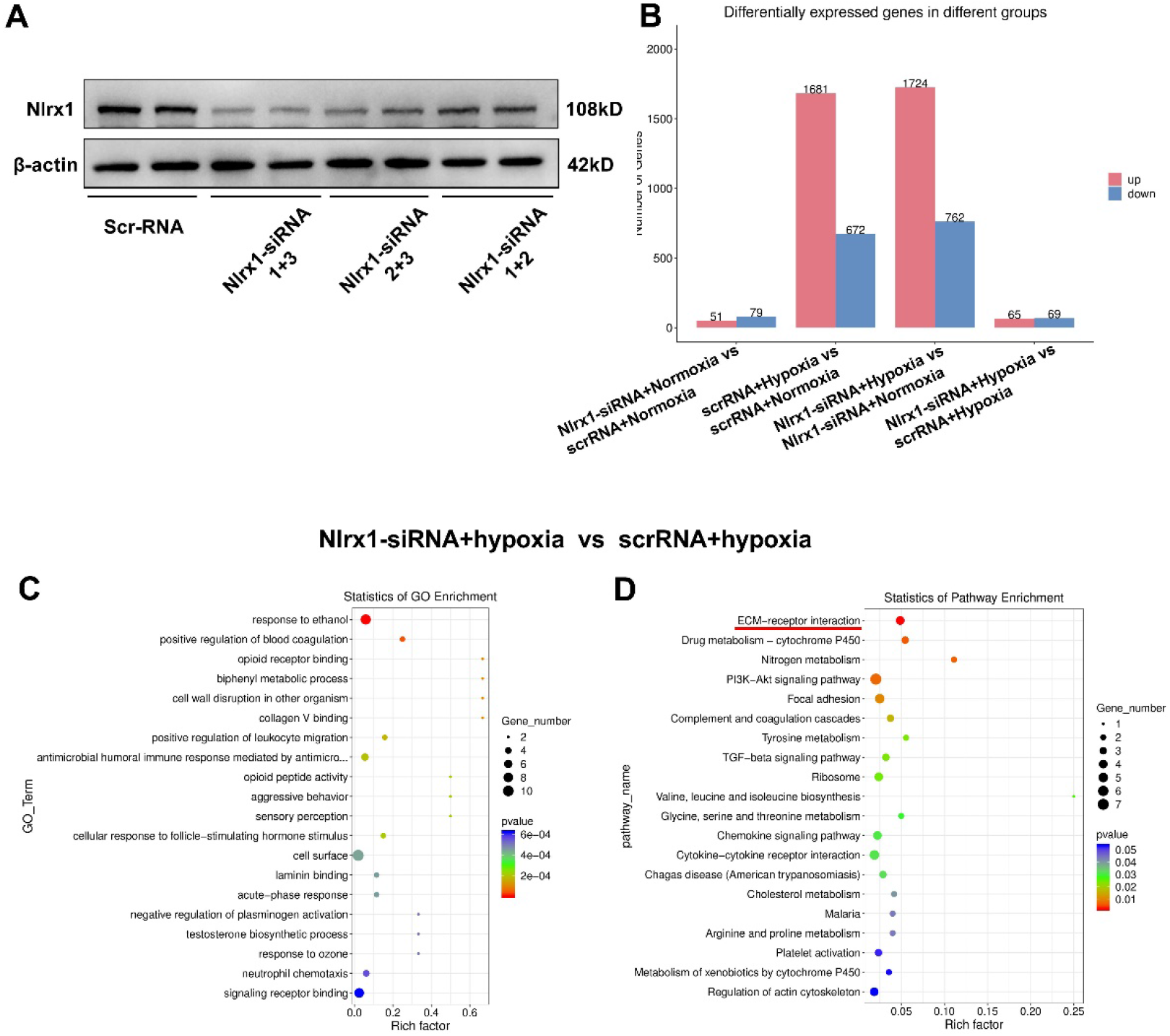
**A**, Representative immunoblot showing knockdown efficacy of Nlrx1. NRVMs were transfected with either Nlrx1-siRNA or scrRNA for 48 hours, and combined use of Nlrx1-targeting siRNA no.1 and siRNA no.3 achieved the most efficacy of knockdown, which was used in the following study to knockdown Nlrx1 expression. **B**, Nlrx1-siRNA or ScrRNA transfected NRVMs were treated by hypoxia and glucose deprivation for 8 hours, then for bulk RNA-seq. Transcriptomic analysis shows the number of DEGs in different groups (>2-fold, padj<0.05). **C-D**, Top 20 of gene ontology (GO) enrichment (**C**) and KEGG pathway enrichment (**D**) of the identified DEGs between Nlrx1-siRNA and ScrRNA transfected NRVMs upon hypoxia and glucose deprivation. DEGs indicates differentially expressed genes; NRVMs, neonatal rat ventricular myocytes; scrRNA, scramble-siRNA.

**Supplementary Figure S18:**
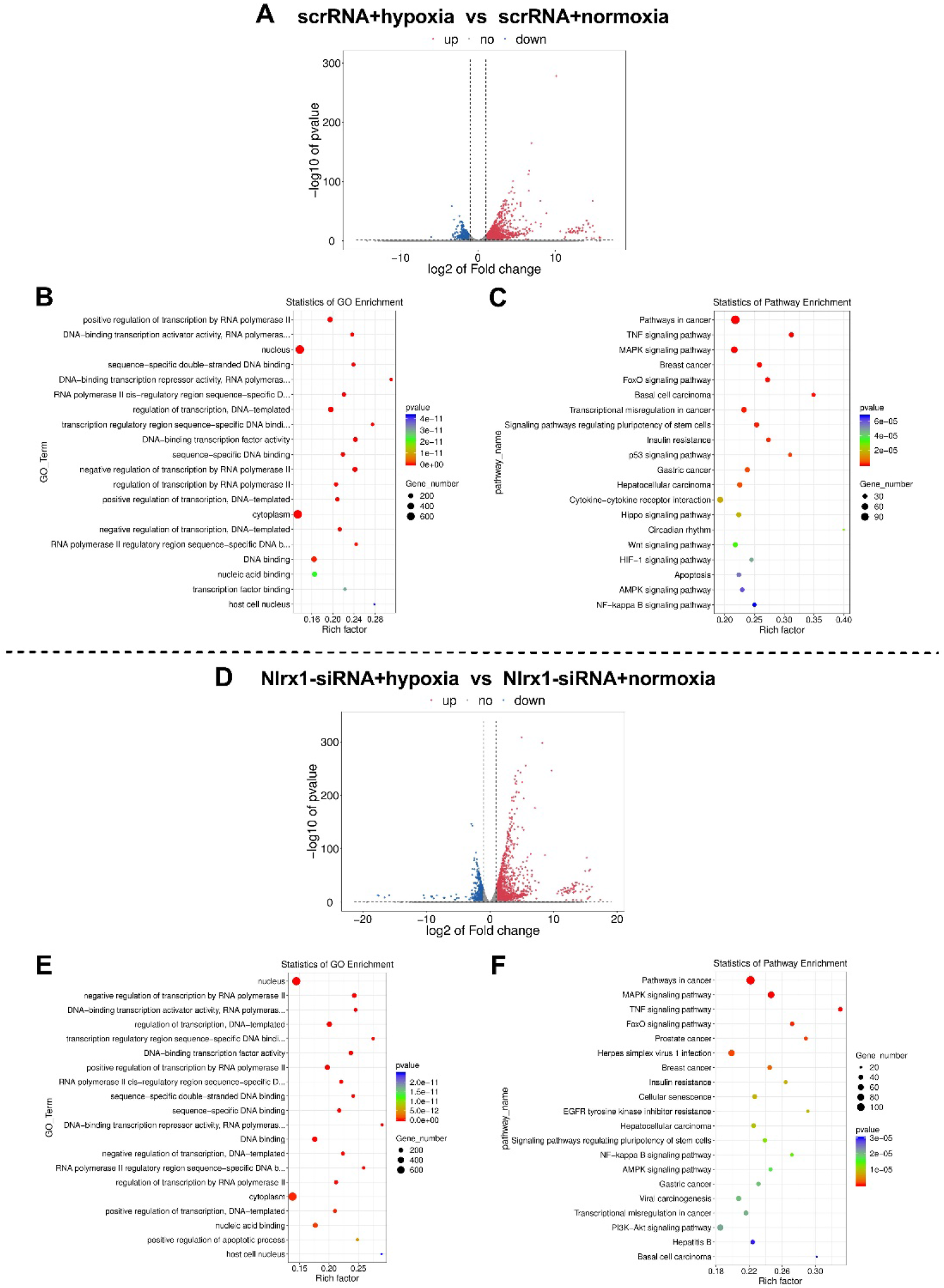
**A**, Volcano plot showing DEGs between hypoxia and normoxia treated NRVMs transfected with scrRNA (>2-fold, padj<0.05). **B-C**, Top 20 of gene ontology (GO) enrichment (**B**) and KEGG pathway enrichment (**C**) of the identified DEGs between hypoxia and normoxia treated NRVMs transfected with scrRNA. **D**, Volcano plot showing DEGs between hypoxia and normoxia treated NRVMs transfected with Nlrx1-siRNA (>2-fold, padj<0.05). **E-F**, Top 20 of gene ontology (GO) enrichment (**E**) and KEGG pathway enrichment (**F**) of the identified DEGs between hypoxia and normoxia treated NRVMs transfected with Nlrx1-siRNA. DEGs indicates differentially expressed genes; NRVMs, neonatal rat ventricular myocytes.

**Supplementary Figure S19:**
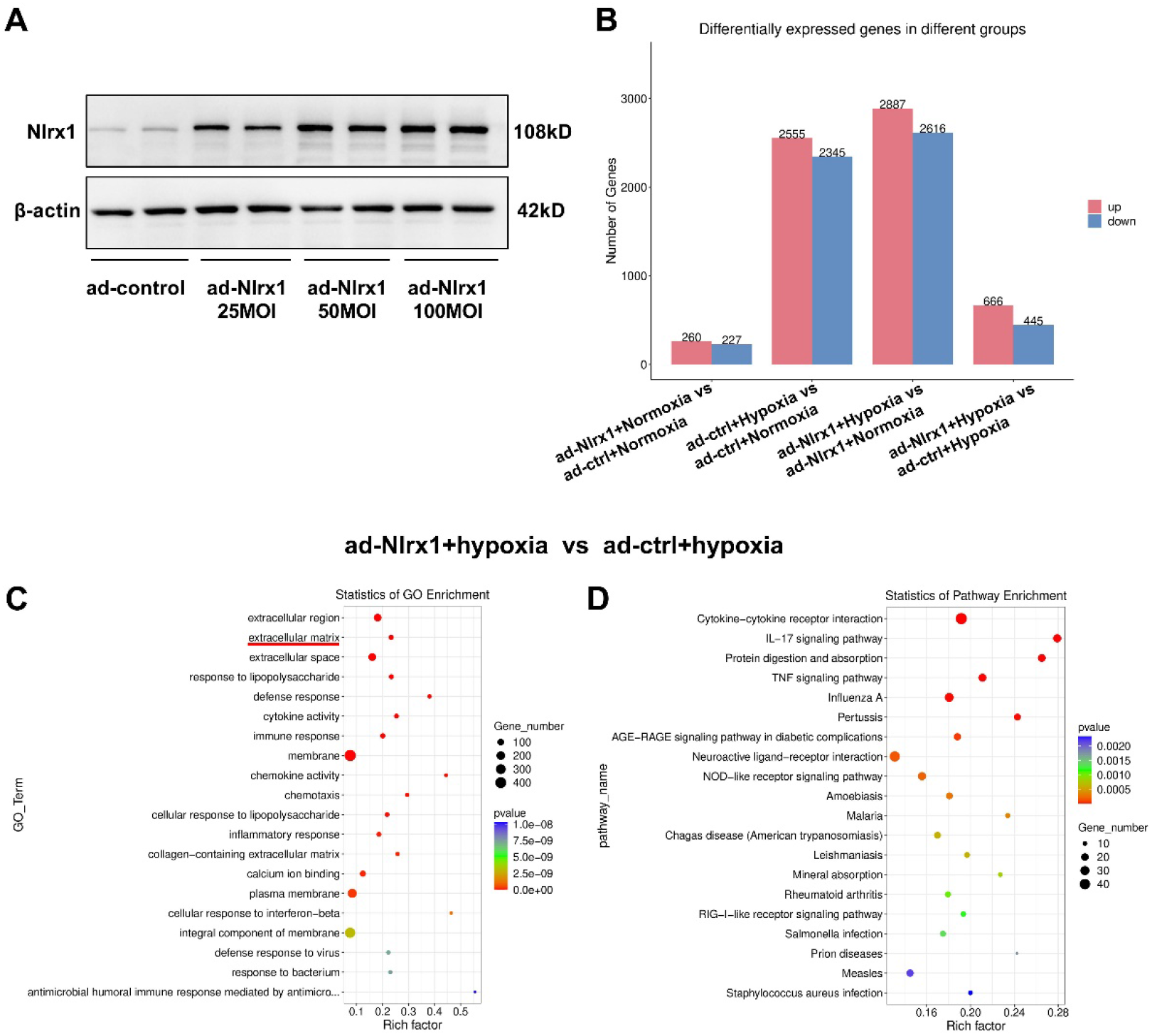
**A**, Representative immunoblot showing overexpression level of Nlrx1. NRVMs were transfected with either ad-Nlrx1 or ad-ctrl of different MOI as indicated for 48 hours, and MOI of 50 was used in the following study to achieve efficient overexpression of Nlrx1 in NRVMs. **B**, ad-Nlrx1 or ad-ctrl transfected NRVMs were treated by hypoxia and glucose deprivation for 8 hours, then for bulk RNA-seq. Transcriptomic analysis shows the number of DEGs in different groups (>2-fold, padj<0.05). **C-D**, Top 20 of gene ontology (GO) enrichment (**C**) and KEGG pathway enrichment (**D**) of the identified DEGs between ad-Nlrx1 and ad-ctrl transfected NRVMs upon hypoxia and glucose deprivation. DEGs indicates differentially expressed genes; NRVMs, neonatal rat ventricular myocytes; ad-Nlrx1, adenovirus-Nlrx1; ad-ctrl, adenovirus-vector.

**Supplementary Figure S20:**
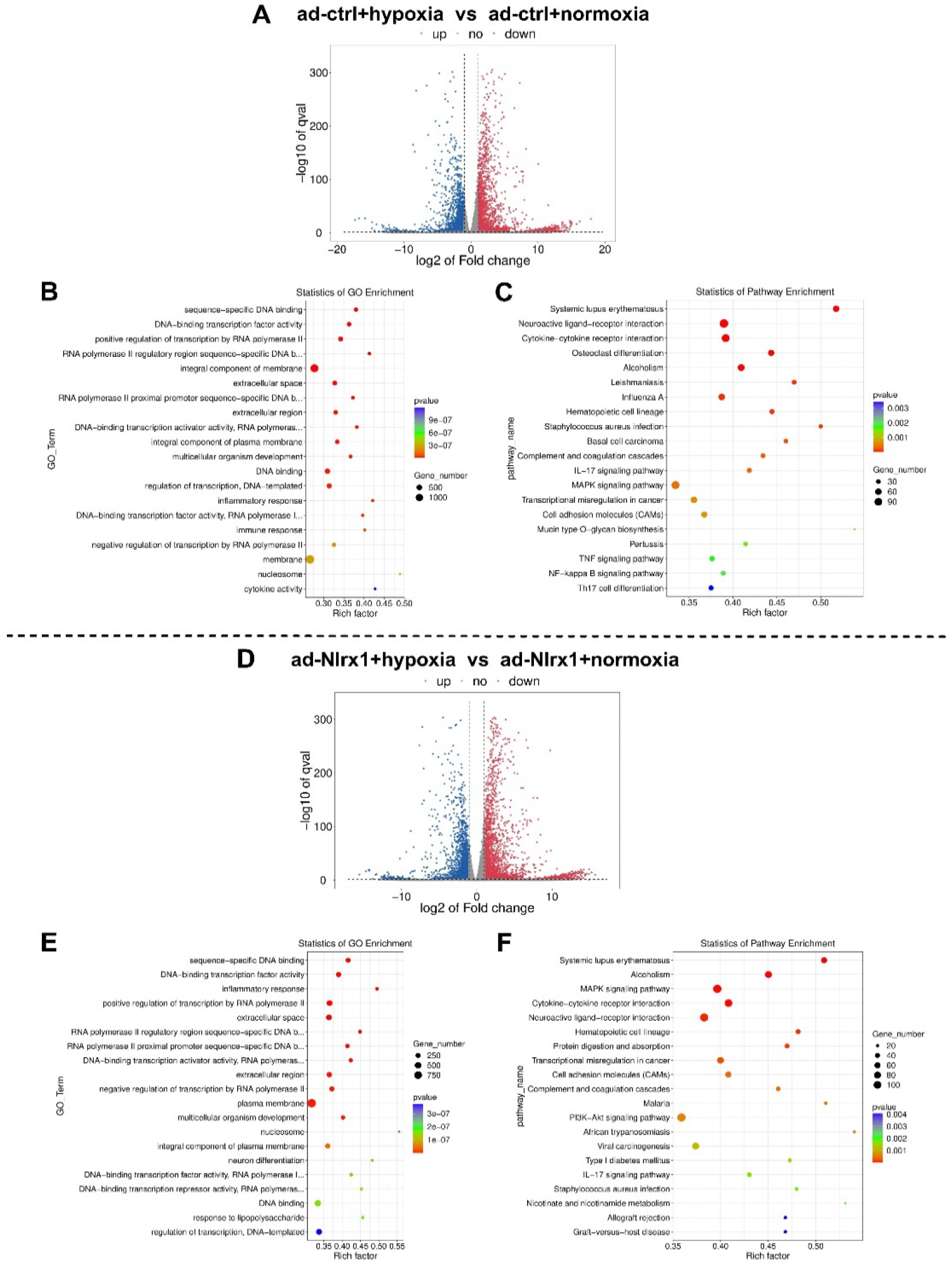
**A**, Volcano plot showing DEGs between hypoxia and normoxia treated NRVMs transfected with ad-ctrl (>2-fold, padj<0.05). **B-C**, Top 20 of gene ontology (GO) enrichment (**B**) and KEGG pathway enrichment (**C**) of the identified DEGs between hypoxia and normoxia treated NRVMs transfected with ad-ctrl. **D**, Volcano plot showing DEGs between hypoxia and normoxia treated Nlrx1-overexpressing NRVMs transfected with ad-Nlrx1 (>2-fold, padj<0.05). **E-F**, Top 20 of gene ontology (GO) enrichment (**E**) and KEGG pathway enrichment (**F**) of the identified DEGs between hypoxia and normoxia treated Nlrx1-overexpressing NRVMs. DEGs indicates differentially expressed genes; NRVMs, neonatal rat ventricular myocytes; ad-Nlrx1, adenovirus-Nlrx1; ad-ctrl, adenovirus-vector.

## Supplemental data of Methods

### Generation of Nlrx1 Knockout Mouse

C57BL/6J wild-type and Nlrx1 knockout (Nlrx1-/-) mouse models were developed by Shanghai Model Organisms Center, Inc. (Shanghai, China). To create the Nlrx1-/- mice, two sgRNAs were designed to delete exons 2-3 of the NLRX1 gene: 5’-CCAAGACATTAGCTGCATGT-3’ and 5’-TTTGGACAGGTCGCGCGTAG-3’. Cas9 mRNA and sgRNAs were transcribed in vitro and injected into zygotes of C57BL/6J mice, which were then transferred to pseudo pregnant recipients. The genotype of the F0 mice was identified and confirmed through PCR and sequencing using the primer pairs F-5’-GCAGGGCAACAGGAAAGAAAGTGG-3’ and R-5’-GGTGGAAGGCCGGGTGGTTAC-3’. Positive F0 mice were chosen and crossed with C57BL/6J mice to obtain F1 heterozygous Nlrx1 knockout mice, which were then confirmed through PCR and sequencing. To obtain the heterozygous Nlrx1-/- mice, F1 heterozygous mice were intercrossed. Male mice aged 8-12 weeks were used for all studies. This study and all animal procedures were performed in accordance with the Guide for the Care and Use of Laboratory Animals (NIH, Bethesda, MD), and were approved by the Animal Care Committee of the Fourth Military Medical University.

### Mouse Model of MI Surgery

Adult male Nlrx1KO and WT mice at the age of 10–14 weeks were subjected to permanent ligation of the left anterior descending (LAD) coronary artery as described previously^1^. Briefly, mice were anesthetized with pentobarbital sodium (50 mg/kg) and then intubated using a 20-gauge intravenous catheter with a blunt end and ventilated with air with a small-animal respirator (tidal volume, 0.3 ml; rate, 150 breaths/ minute; Harvard Apparatus, USA). The chest wall was shaved, and a left thoracotomy was performed in the fourth intercostal space. The left ventricle and the LAD coronary artery were visualized using a microscope and the LAD artery was permanently ligated with a 7-0 monofilament suture at 1-2 mm distal to its emergence from under the left atrium. Infarction was confirmed by the presence of myocardial blanching in the ischemic area and ST segment elevation on the electrocardiogram. The thoracotomy was closed with 5-0 silk sutures. The endotracheal tube was removed once spontaneous respiration resumed, and animals were placed on a warm pad maintained at 37 ℃ until they were completely awake. Sham-operated animals underwent the same procedure without coronary artery ligation. Mice that did not survive the recovery from anesthesia and mice that died within 24 hours after surgery were excluded from the experiment.

### Echocardiography

Serial transthoracic echocardiographic measurements at baseline, 1 week after MI surgery was performed with a VEVO 2100 platform (Visual Sonics, Toronto, Canada) as described previously. Mice were anesthetized by inhalation of isoflurane (1–2%) in a 100% oxygen mix, and heart rate was maintained at approximately 400-500 bpm in all mice during the echocardiographic examination to minimize data deviation. M-mode tracings were recorded through the anterior and posterior LV walls at the papillary muscle level to measure LV end-diastolic dimension (LVEDD) and LV end-systolic dimension (LVESD). The ejection fraction was calculated with the formula: [(LVEDV−LVESV)/LVEDV]×100, where EDV is end-diastolic volume and ESV is end-systolic volume. All the echocardiographic images were analyzed using Vevo 2100 software. The measurements represented the average of 6 selected cardiac cycles from at least 2 separate scans performed in random-blind manner.

### Histological analysis

Hearts were arrested at diastole by injection of phosphate buffer saline. Heart tissues were harvested and immediately immersed in 4% paraformaldehyde (pH 7.4). Following embedded in paraffin, and sliced into 5-µm sections. Sections were deparaffinized and rehydrated followed by staining with hematoxylin and eosin (HE dye solution kit, Servicebio, G1003), Masson trichrome (Masson dye solution kit, Servicebio, G1006), and picrosirius red (Sirius Red solution set, Servicebio, G1018) to determine the collagen volume fraction and morphological effects. Staining was performed following manufacturer’s standard procedures. Images were acquired using Pannoramic DESK (3D HISTECH). Histological figures showing large longitudinal mice heart sections were captured using CaseViewer software (3D HISTECH). 5 randomly chosen high-power fields (× 200) in each section were captured and analyzed via Image J. Obtained analysis of results from all slides in the same heart were averaged and counted as n=1.

### Flow cytometric sorting of infarct macrophages

Cardiac CD11b^+^Ly6G^-^CD9^+^ macrophages of infarcted hearts were harvested through flow cytometric sorting. Briefly, infarcted heart tissues including infarct zone and border zone were minced and placed into a cocktail of GEXSCOPE® sCelLiVE Tissue Dissociation Solution kit (Singleron Biotechnologies, Nanjing, China), and shaken at 37°C for 15 minutes twice for tissue dissociation. Then cells were passed through a 70-μm nylon mesh (Corning Falcon, 352350) and collected for centrifugation (10 minutes, 300 g, 4°C). The cell pellet was resuspended in 2 mL of sterile red blood cell lysis buffer (Biolegend, 420302) for 10 min to remove the erythrocytes. Cell preparations were stained in 200 μL of cell staining buffer at 4°C for 30 min with the following antibodies: anti-mouse/human CD11b-FITC (clone M1/70, Biolegend, catalog no. 101206), anti-mouse CD9-APC (clone MZ3, Biolegend, catalog no. 124812), anti-mouse Ly-6G-Pacific Blue™ (clone 1A8, Biolegend, catalog no. 127612), then washed, centrifuged and resuspended for FACS sorting (BD FACSAria™II). Dosages of all antibodies were determined per 1·10^7^ cells. FlowJo software (San Carlos, CA) was applied to analyze examined results. Total RNA was extracted from sorted macrophages using TRIzol Reagent (Invitrogen) for RNA-seq study.

### Neonatal rat ventricular myocyte (NRVMs) culture and in vitro hypoxia assay

NRVMs were prepared by enzymatic digestion from 1- to 2-day-old Sprague–Dawley neonatal rats as previously described. Briefly, ventricles of hearts were minced into ≈1 mm pieces and digested with 1mg/mL Type I collagenase (Gibco, 17100017) at 37°C for 15 minutes, and the resultant cell suspensions were seeded into DMEM containing 10% (v/v) fetal bovine serum (FBS) for 1.5 hours. Afterwards, fibroblasts were removed, and cardiomyocytes were reseeded into fresh DMEM supplemented with 10% FBS, 2 mM L -glutamine, and penicillin–streptomycin. To simulate hypoxia in vitro, NRVMs were placed in the hypoxia chamber (95% N_2_ + 5% CO_2_) in glucose-free DMEM medium for 9 hours for RNA-seq. Cardiomyocytes were harvested for further experiments at different time points.

### Nlrx1-overexpression in NRVMs by Adenovirus infection

Adenoviral infection of NRVMs was employed to overexpress Nlrx1. At 24 h after plating, NRVMs were transfected with adenoviruses expressing Nlrx1 (Ad-Nlrx1) or adenoviruses with an empty vector (Ad-control) at varying multiplicities of infection (MOIs) (Hanbio Technology, Shanghai, China) in DMEM medium. After 24 hours of adenoviral infection, the medium was replaced with virus-free SFM medium, and cells were cultured for additional 12 hours prior to subsequent experiments. Levels of expressed proteins were determined by Western blot analysis.

### SiRNA mediated knockdown of Nlrx1 in NRVMs

Pre-designed SiRNA targeting Nlrx1 and SiRNA control were obtained from Hanbio Technology (Shanghai, China) and sequence of SiRNA were listed in Table 1. SiRNA transfection was performed using LipofectamineTM RNAiMAX reagent (Invitrogen, 13778150) according to the manufacturer’s instructions. Briefly, 6 x10^5^ cells were transfected in 2ml of Opti-MEM containing 7µl of Lipofectamine RNAiMAX and 100 nmol of SiRNA. The efficacy of Nlrx1 knockdown was determined by Western blot analysis.

**Table 1.**
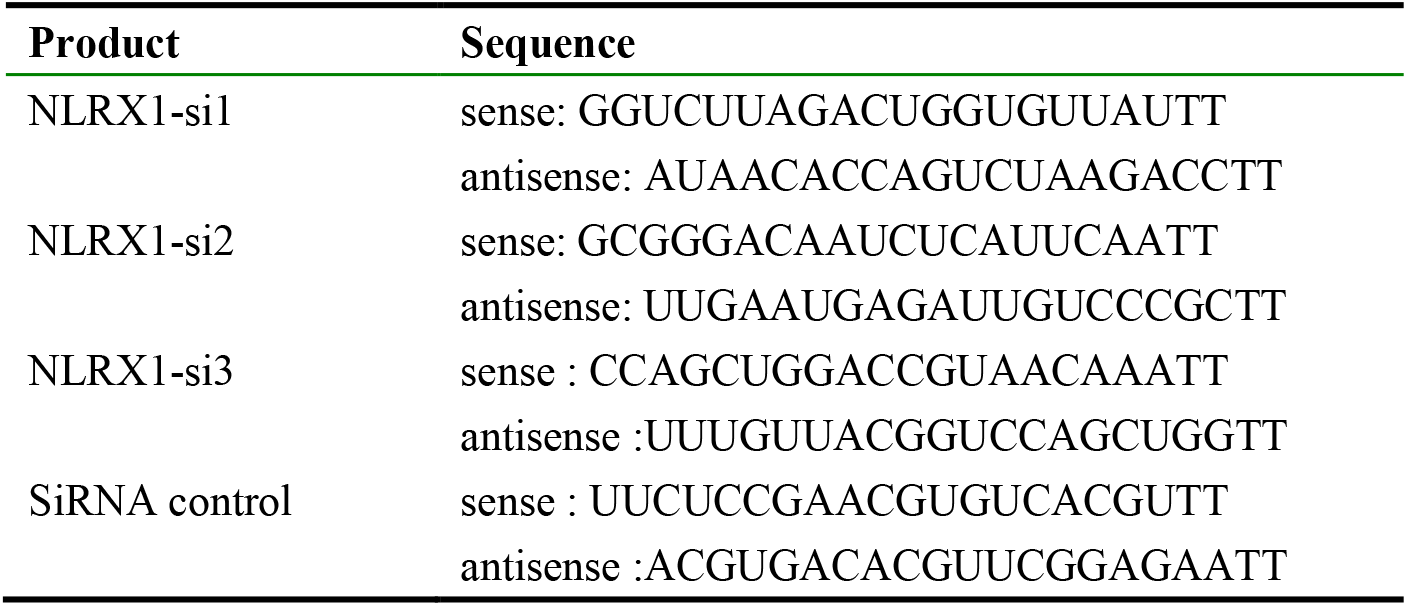

### Bone marrow-derived macrophages (BMDM) and in vitro macrophages stimulation

Bone marrow cells were collected as previously reported^2^. Briefly, femurs and tibiae of wild type mice were carefully flushed with pre-cooled PBS containing 2% FBS by inserting a syringe into the medullary cavity at one end of the bone, and the bone marrow was collected and filtered through a 70 µm nylon mesh, followed centrifugation at 1000 rpm for 10 min and resuspended in RBC Lysis buffer, then cultured in DMEM medium (Gibco) containing macrophage colony-stimulating factor (M-CSF 20ng/mL, Sino Biological, Catalog no. 51112-M08H) for 7 days. The medium was changed every 3 days.

Conditioned medium from neonatal rat ventricular myocytes (NRVMs) upon hypoxia stimulation and glucose deprivation for 24 hours, were used to stimulate BMDMs for 24 hours. BMDMs were stimulated with recombinant human CCN2 (MCE, Catalog no. HY-P72153) for 24 hours with or without conditioned medium from neonatal rat ventricular myocytes (NRVMs) upon hypoxia. Total RNA was extracted and gene expression was determined by quantitative RT-PCR.

### RNA isolation and quantitative real-time polymerase chain reaction (PCR)

Total RNA was extracted using the Trizol Substitute (Solarbio Beijing China) according to the manufacturer’s instruction. RNA concentration was quantified and checked for purity (OD 260/280) using a Nano-Drop-1000 spectrophotometer (ThermoFisher). Reverse transcription of equal RNA content was performed using the Goldenstar™ RT6 cDNA Synthesis Kit (Tsingke Biotechnology TSK314S), and real-time quantitative PCR was conducted using 2×TSINGKE® Master qPCR Mix (SYBR Green I) (Tsingke TSE201) in accordance with the manufacturer’s protocol. The primer pairs were designed using Primer Premier 5.0 software and sequences were listed in Table 2. The gene expression levels were normalized to the reference gene Actin and the data were reported as 2^-ΔCt^ values± SEM.

**Table 2.**
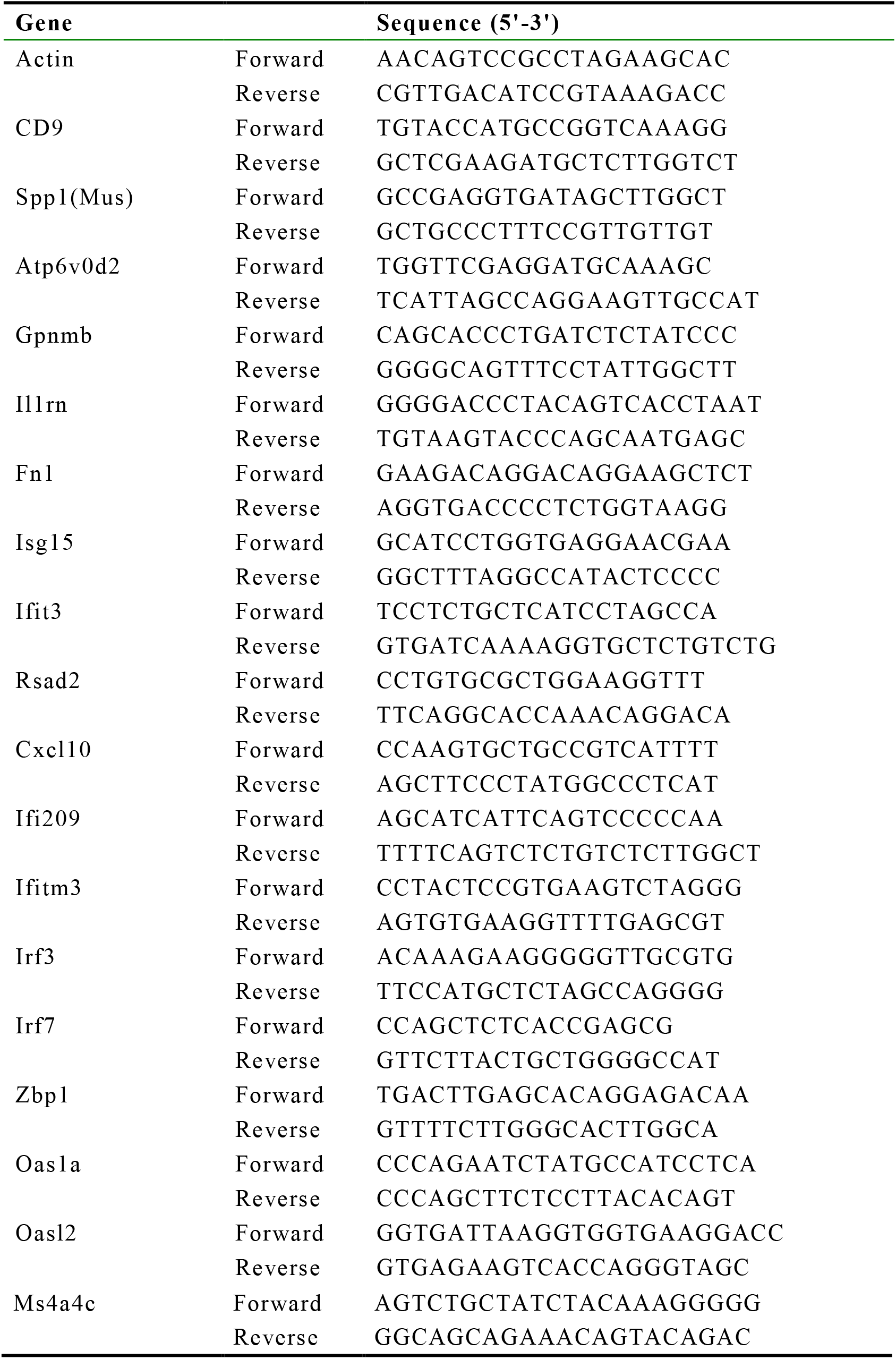

### Western blot analysis

Total protein was extracted using RIPA buffer containing protease inhibitor cocktail. Equal amounts of protein were mixed with sample loading buffer and under reducing conditions separated on SDS-polyacrylamide gel. Proteins were electro-transferred on PVDF membranes (0.2 µ m, Millipore). Blots were incubated with following primary antibodies overnight at 4 °C: anti-NLRX1 antibody (Immunoway, Cat No.YT5802), anti-CTGF antibody (Abcam, Clone: EPR20728, ab209780), β-Actin (Proteintech, Cat No.81115-1-RR). The secondary antibodies were horseradish peroxidase-conjugated goat antirabbit secondary antibody (A0208; Beyotime Biotechnology). The bands were detected using a chemiluminescence assay (ECL, Millipore).

### Tissue dissociation and single-cell suspension preparation

Infarcted heart tissues (infarct zone and border zone) were isolated on day 3 after MI, washed with Hanks Balanced Salt Solution (HBSS) for three times, minced into small pieces, and digested in 2 ml of GEXSCOPE Tissue Dissociation Solution (Singleron Biotechnologies) following manufacturer’s instructions. Briefly, the heart tissues were digested at 37 °C for 15min with continuous agitation. A 40-micron sterile strainer (Corning) was used to separate cells from impurities after digestion. The cells were then centrifuged at 300 ×g 4 °C for 5min and cell pellets were resuspended in 1ml PBS (HyClone). Cell suspensions were counted with TC20 automated cell counter (Bio-Rad) to determine cell concentration and viability.

### Single-cell RNA-sequencing library preparation

The concentration of single-cell suspension was adjusted to 1× 10^5^ cells/mL in PBS. Single-cell suspension was then loaded onto a microfluidic chip (GEXSCOPE Single Cell RNA-seq Kit, Singleron Biotechnologies) and scRNA-seq libraries were constructed according to the manufacturer’s instructions (Singleron Biotechnologies). The resulting scRNA-seq libraries were sequenced on an Illumina novaseq 6000 with 150 bp paired end reads.

### Generation of single-cell gene expression matrices

Raw reads were processed to generate gene expression matrices by CeleScope (https://github.com/singleron-RD/CeleScope) pipeline. Briefly, raw reads were first processed with CeleScope to remove low quality reads with Cutadapt v1.17 to trim poly-A tail and adapter sequences^3^. Cell barcode and UMI were extracted. After that, we used STAR v2.6.1a^4^ to map reads to the reference genome GRCm38 (ensembl version 92 annotation). UMI counts and gene counts of each cell were acquired with featureCounts v2.0.1 software^5^, and used to generate expression matrix files for subsequent analysis.

### Quality control, cell-type clustering, and major cell-type identification

Cells with gene count less than 200 or with top 2% gene count were removed. Furthermore, cells with top 2% UMI count and mitochondria content higher than 50% were discarded. Finally, 23,805 cells including 12,180 cells from 3 WT infarcted hearts and 11,625 cells from 3 Nlrx1KO infarcted hearts, were obtained for the downstream analysis. We obtained 1858 genes and 5615 UMIs per cell on average. We used functions from Seurat v3.1.2 for dimension-reduction and clustering^6^. Then we used NormalizeData and ScaleData functions to normalize and scale all gene expression, and selected the top 2000 variable genes with FindVariableFeautres function for PCA analysis. Using the top 20 principle components, we separated cells into 37 cell clusters. To assign one of the 11 major cell types to each cluster, we scored each cluster by the normalized expressions of the following canonical markers: Endothelial cells (Cdh5, Pecam1, Kdr), Fibroblasts (Col1a1, Col1a2, Dcn, Lum), Macrophages (CD68, CD163, Lyz2), Monocytes (Ly6c, Lyz2, Sell), Neutrophils (CSF3R, S100a8, Cxcr2), Dendritic cells (CD209a, H2-Ab1, Clec10a), Pericytes (Rgs5, Mcam, Pdgfrb), Smooth muscle cells (Acta2, Tagln, Mylk), T cells (CD2, CD3d, Trac), B cells (CD79a, CD79b, Ms4a1). The highest scored cell type was assigned to each cluster. The clusters assigned to the same cell type were lumped together for the following analysis. The final results were manually examined to ensure the correctness of the results and visualized by Uniform Manifold Approximation and Projection (UMAP).

### Differentially expressed genes (DEGs) analysis

To identify differentially expressed genes (DEGs), we used the Seurat FindMarkers function based on Wilcox likelihood-ratio test with default parameters, and selected the genes expressed in more than 10% of the cells in a cluster and with an average log (Fold Change) value greater than 0.25 as DEGs. For the cell type annotation of each cluster, we combined the expression of canonical markers found in the DEGs with knowledge from literatures, and displayed the expression of markers of each cell type with heatmaps/dot plots/violin plots that were generated with Seurat DoHeatmap/DotPlot/Vlnplot function. Doublet cells were identified as expressing markers for different cell types, and removed manually.

### Pathway enrichment analysis

To investigate the potential functions of DEGs, the Gene Ontology (GO) and Kyoto Encyclopedia of Genes and Genomes (KEGG) analysis were used with the “clusterProfiler” R package 3.16.1^7^. Pathways with p-adj value less than 0.05 were considered as significantly enriched. Gene Ontology gene sets including molecular function (MF), biological process (BP), and cellular component (CC) categories were used as reference.

### Trajectory analysis

Cell differentiation trajectory was reconstructed with Monocle2^8^. Highly-variable genes (HVGs) were used to sort cells in order of spatial-temporal differentiation. We used DDRTree to perform FindVairableFeatures and dimension-reduction. Finally, the trajectory was visualized by plot_cell_trajectory function. Next, CytoTRACE^9^ (a computational method that predicts the differentiation state of cells from single-cell RNA-sequencing data using gene Counts and Expression) was used to predict the differentiation potential of monocyte subpopulations.

### RNA velocity

For RNA velocity, BAM file and reference genome GRCm38 were used in the analysis with velocyto v0.2.3 and scVelo v0.17.17 in python with default parameters^10,11^. The result was projected to the UMAP plot from Seurat clustering analysis for visualization consistency.

### Library preparation for transcriptome sequencing

RNA samples from macrophages of infarcted hearts or NRVMs were sent to LC-Bio Technology (Hangzhou, China) to construct cDNA libraries by using NEBNext® Ultra™ RNA Library Prep Kit for Illumina® (NEB, Ipswich, MA, USA). In brief, the process of library construction consisted of i) mRNA purification and enrichment from total RNA (5μg) using oligo(dt)-attached magnetic beads (Dynabeads Oligo (dT), Thermo Fisher, CA, USA), ii) fragmentation of purified mRNA using divalent cations exposed to elevated temperatures in Magnesium RNA Fragmentation Module (NEB, cat.e6150, USA), iii) double stranded cDNA synthesis, using Rnase H-reverse transcriptase (first strand) and DNA polymerase I, dNTP and RNase H (second strand); iv) terminal cDNA ends reparation by exonucleases/polymerases; and poly adenylation of the 3’ ends of the DNA fragments, v) sequencing adaptors ligation, and vi) cDNA fragments size selection (150-200 bp in length) which underwent PCR. PCR was performed using Phusion High-Fidelity DNA Polymerase, universal PCR primers and Index (X) Primer. PCR products were purified (AMPure XP system), and the library concentration (>2 nM) and quality were assessed using a Bioanalyzer 2100 system (Agilent, Santa Clara, CA, USA).

### RNA-sequencing: quality analysis, mapping and assembly

The library preparations were sequenced on Illumina Novaseq 6000 devices (LC-Bio Technology CO., Ltd., Hangzhou, China), generating 2×150 bp paired-end reads. Adapter, poly-N and low-quality reads from the raw data were excluded to purify the data analysis. Quality Phred Scores, Q20%, Q30% and GC contents of the clean data were calculated, showing high accuracy of reads (>99%, with 0.02 error rate). Filtered reads were aligned to the C57BL/6J reference genome (December 2011 release of the mouse musculus reference genome (mm10; GRCm38.p6) from Ensembl) using HISAT2 algorithm alignment program. Mapped reads were assembled using stringTie.

### Gene expression, differential expression, enrichment and co-expression analysis

HTSeq software (versition: hisat2-2.2.1) was used to count the number of reads mapped to each gene. Read count of fragments per kilobase of transcript sequence per millions base pairs sequenced (FPKM) was used to calculate gene expression level, which considered the effects of both sequencing depth and gene length. Readcount obtained from the gene expression analysis was used for differential expression analysis. Cluster differential expression analysis for every gene in different conditions was performed using the DESeq2 R software package (v.1.10.1). Genes with an average log (Fold Change) value greater than 2 and adjusted P-value ≤ 0.05 were considered to be differentially expressed. List of genes were ranked by differential gene expression as log2 (fold change) between each comparison group. Positive values were upregulated genes, whereas negative values were downregulated genes. All RNA-seq processed data have been deposited in NCBI’s Gene Expression Omnibus (GEO) and are accessible through GEO SuperSeries accession number.

### Statistical analysis

For all analyses, normal distribution was tested using the Shapiro-Wilk normality test. For comparisons of two groups, unpaired two-tailed Student’s t test using Welch’s correction for unequal variances was performed. For comparisons of multiple groups, one-way ANOVA was performed followed by Tukey’s multiple comparison test. Survival analysis was performed using the Kaplan-Meier method. Mortality was compared using the log rank test. Data are expressed as means ± SEM. Statistical significance was set at P < 0.05.

